# Mapping the Macrostructure and Microstructure of the in vivo Human Hippocampus using Diffusion MRI

**DOI:** 10.1101/2022.07.29.502031

**Authors:** Bradley G. Karat, Jordan DeKraker, Uzair Hussain, Stefan Köhler, Ali R. Khan

## Abstract

The hippocampus is classically divided into mesoscopic subfields which contain varying microstructure that contribute to their unique functional roles. It has been challenging to characterize this microstructure with current MR based neuroimaging techniques. In this work, we used diffusion MRI and a novel surface-based approach in the hippocampus which revealed distinct microstructural distributions of neurite density and dispersion, T1w/T2w ratio as a proxy for myelin content, fractional anisotropy, and mean diffusivity. We used the Neurite Orientation Dispersion and Density Imaging (NODDI) model optimized for gray matter diffusivity to characterize neurite density and dispersion. We found that neurite dispersion was highest in the Cornu Ammonis (CA) 1 and subiculum subfields which likely captures the large heterogeneity of tangential and radial fibers, such as the Schaffer collaterals, perforant path, and pyramidal neurons. Neurite density and T1w/T2w were highest in the subiculum and CA3 and lowest in CA1, which may reflect known myeloarchitecture differences between these subfields. Using a simple logistic regression model, we showed that neurite density, dispersion, and T1w/T2w measures provided good separability across the subfields, suggesting that they may be sensitive to the known variability in subfield cyto- and myeloarchitecture. We report macrostructural measures of gyrification, thickness, and curvature that were in line with ex vivo descriptions of hippocampal anatomy. We employed a multivariate orthogonal projective non-negative matrix factorization (OPNNMF) approach to capture co-varying regions of macro- and microstructure across the hippocampus. The clusters were highly variable along the medial-lateral (proximal-distal) direction, likely reflecting known differences in morphology, cytoarchitectonic profiles, and connectivity. Finally, we show that by examining the main direction of diffusion relative to canonical hippocampal axes, we could identify regions with stereotyped microstructural orientations that may map onto specific fiber pathways, such as the Schaffer collaterals, perforant path, fimbria, and alveus. These results highlight the value of combining in vivo diffusion MRI with computational approaches for capturing hippocampal microstructure, which may provide useful features for understanding cognition and for diagnosis of disease states.

## 1. Introduction

The hippocampus is classically divided into structurally distinct mesoscopic subfields according to differences in cyto-, myelo-, and chemoarchitecture (Duvernoy et al., 2013; Ding & Van Hoesen, 2015). The neuronal circuitry that compose the microstructure of the hippocampus exist within and across these subfields. For example, the pyramidal neurons that exist within the Cornu Ammonis (CA) and subiculum subfields have apical and basal dendrites which project across the laminae, while their axons project to the alveus, a white matter bundle adjoining the hippocampus. The trisynaptic pathway is the major circuitry component which connects the subfields of the hippocampus. The entorhinal cortex connects to the dentate gyrus (DG) and other subfields through the myelinated perforant path. The DG then projects to the pyramidal neurons of CA3 through the mossy fibers, which then project to CA1 through the largely unmyelinated Schaffer collaterals. Finally, CA1 projects to the subiculum and back to the entorhinal cortex as a large hippocampal efferent. Hippocampal microstructure is key in producing unique cognitive functions such as memory formation and storage and spatial navigation among others (Voss et al., 2017; Goodroe et al., 2018; Horner et al., 2015). Furthermore, the hippocampus is typically one of the earliest aberrant structures in many disease states, where specific microstructural properties are differentially afflicted or spared (Moodley & Chan, 2014; Dhikav & Anand, 2012; Small et al., 2011). While much work has addressed volumetric characterization of the hippocampus and its subfields, understanding hippocampal microstructure can provide key insights into its complex cognitive functions as well as its early deterioration in disease.

Diffusion magnetic resonance imaging (dMRI) is a particular technique which holds promise in probing the hippocampal circuitry by sensitizing the measured MRI signal to the movement of water molecules, which diffuse more readily parallel to microstructure. Several models have been proposed that attribute measures of the dMRI signal to compartments which have varying diffusivity environments (Assaf et al., 2008; Assaf & Basser, 2005; Zhang et al., 2012). One of the earliest and most widely used models proposed by Basser et al. (1994) is diffusion tensor imaging (DTI). DTI estimates quantitative parameters such as fractional anisotropy (FA - a measure of the anisotropy of diffusion), mean diffusivity (MD – magnitude of diffusion), and the ellipsoidal orientation of the diffusion process. However, DTI has some notable limitations. At increasing b-values (approximately greater than 1000-1500 *s*/*mm*^2^) there is contribution from multiple compartments with varying diffusivities (such as restricted intra-cellular water), which goes beyond the monoexponential signal modelled by DTI (Assaf & Cohen. 2000). As well, regions of crossing fibers result in planar DTI ellipsoids with understated FA values (Campbell et al., 2005). Furthermore, DTI measures are sensitive to multiple microstructural properties at the same time, decreasing their specificity (Pierpaoli et al., 1996). Other models aim to utilize increasingly complex diffusion acquisitions to relate the diffusion signal attenuation to varying sets of biophysically motivated compartments.

One popular compartment model is Neurite Orientation Dispersion and Density Imaging (NODDI), which aims to provide a biophysical interpretation of the diffusion signal (Zhang et al., 2012). NODDI assumes that three microstructural environments consisting of an intra-cellular, extra-cellular, and cerebrospinal fluid (CSF) compartment contribute to the diffusion signal. The intra-cellular compartment is modeled as a set of infinitely anisotropic sticks (diffusion can only be parallel to the main orientation of the stick), while the extra-cellular compartment is modeled as a zeppelin (or cylindrically symmetric tensor) with hindered diffusion perpendicular to its main axis. The CSF compartment is modeled as a sphere with gaussian isotropic diffusion. Diffusion is assumed to be contained separately within each compartment, where the resulting signal is the sum from all compartments. NODDI aims to overcome the limitations of DTI by providing microstructural scalars such as the neurite density index (NDI) and orientation dispersion index (ODI) which are sensitive to fiber crossings and are biophysically grounded (Zhang et al., 2012). NDI is meant to represent the volume fraction of the intra-cellular compartment, which is believed to correspond to the density of dendrites, axons, and other “stick” like processes in a voxel. ODI is meant to capture the essence of the orientation distribution of the diffusion signal, where more complex or disperse microstructural configurations correspond to a higher ODI value.

Contemporary work has attempted to examine hippocampal microstructure with both DTI and NODDI. Some such studies have found age-related changes of hippocampal microstructure by averaging NODDI measures within subfields (Radhakrishnan et al., 2020) while others have shown regionally specific changes using DTI (Yassa et al., 2010). Recent work has also begun to use measures derived from structural MRI to interrogate microstructural characteristics, such as the ratio of T1-weighted over T2-weighted signal as a correlate of myelin (Glasser & Van Essen, 2011; Glasser et al., 2014). These measures may be useful to capture the myelinated intra-hippocampal circuitry like the perforant path. A recent study investigated the variation of DTI and intra-cortical myelin through the ratio of T1w/T2w images across the hippocampus using non-negative matrix factorization, however, they did not make quantitative comparisons of microstructure within and across the subfields (Patel et al., 2020). As well, they note the limited specificity nature of the DTI and T1w/T2w metrics investigated. A recent study examined the distribution of NODDI metrics and cortical thickness across the entire cerebral cortex including the hippocampus by averaging metrics across all subjects within each cortical parcel (Fukutomi et al., 2018). Thus, they only examined coarse-grained averages across the entire hippocampal volume. The spatial distributions of NODDI and DTI measures have not been extensively investigated within the hippocampal subfields and across its longitudinal axis.

The orientation and trajectory of the hippocampal circuitry including the trisynaptic circuit has been probed previously using tractography and polarized light imaging (PLI). Ex vivo work has indeed resolved major parts of the hippocampal circuitry using dMRI tractography (Beaujoin et al., 2018) and PLI (Zeineh et al., 2017) in a small number of samples. While these studies serve as a close to ground-truth reference for the orientation of hippocampal circuitry, a difficult step has been recapitulation of this circuitry in vivo. Some in vivo work has attempted to use DTI to capture parts of the trisynaptic circuit such as the perforant path (Yassa et al., 2010) or the whole hippocampal circuitry (Zeineh et al., 2012). However, it is unclear whether the found trajectories are anatomically valid. Furthermore, at lower resolutions, tracts can be spurious requiring complex acquisition and correction schemes, and since acquisitions can vary across studies, tractography practically always requires separate optimization of its parameters (Zeineh et al., 2012). Thus, understanding hippocampal microstructure in vivo may benefit from a simpler characterization of the orientation of hippocampal circuitry with reference to metrics derived from common models like NODDI within and across the subfields.

In the current study, we examined the spatial distribution of NODDI and DTI metrics, T1w/T2w ratio, and macrostructural features of thickness, gyrification, and curvature across the hippocampus using high-resolution in vivo human connectome project (HCP) data (Van Essen et al., 2013), something that has not been extensively investigated. We aimed to examine if the microstructural metrics systematically vary across the cytoarchitectonic defined subfields. Furthermore, we used Orthogonal Projective Non-Negative Matrix Factorization (OPNNMF) as a multivariate approach to capture regions of the hippocampus where these metrics co-vary. We aimed to consider the current OPNNMF representation of disparate macro- and microstructural metrics under the framework of previous research which has suggested modes of hippocampal organization along its medial-lateral (across subfields) and anterior-posterior (longitudinal) axes (Genon et al., 2021; Robinson et al., 2015; Zhong et al., 2019; Cheng et al., 2020; Plachti et al., 2019; Plachti et al., 2020; Patel et al., 2020, DeKraker et al., 2020). While most hippocampus representations use voxel-based approaches, we utilized a novel surface-based approach called *HippUnfold* (DeKraker et al., 2018; DeKraker et al., 2022). Much like in the neocortex, surface-based methods are better suited to account for interindividual differences in tissue curvature and digitation than voxel-based approaches (DeKraker et al., 2022; DeKraker et al., 2021). Aligning hippocampi on a 2D surface preserves topology and the known contiguity of subfields, allowing for improved anatomical detail to be captured. Finally, hippocampal gray matter shows a laminar distribution similar to that of other cortical areas with large radial and tangential neurite components, although the highly curved structure of the hippocampus is reflected in the complexity of its neurite orientations. Importantly, these neurite orientations tend to be highly aligned along one of the axes of the hippocampus that span the anterior-posterior (AP longitudinal), proximal-distal (PD - across subfields), or inner-outer (across laminae) directions (Figure 1A and B) (Zeineh et al., 2017; Nieuwenhuys et al., 2008; Duvernoy et al., 2013). Thus, we also aimed to determine if the known stereotyped orientations of microstructure can be elucidated by analyzing the primary direction of diffusion along each of the axes in vivo, as depicted in Figure 1B.

**Figure 1.**
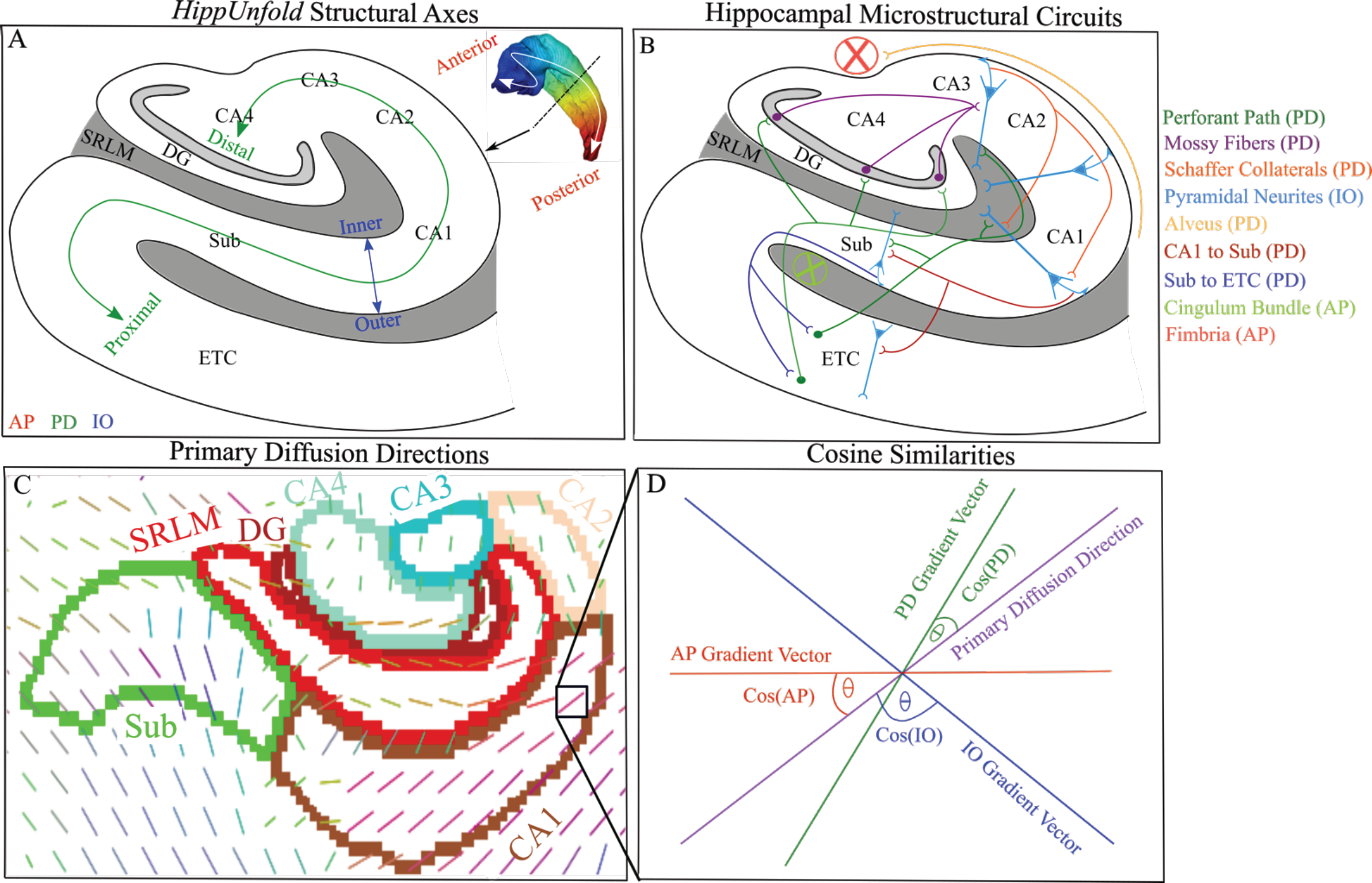
Depiction of hippocampal structural axes, the stereotyped organization of microstructure, and diffusion vectors of the hippocampus. (A) A coronal slice depicting the structural axes of the hippocampus defined as anterior-posterior (AP), proximal-distal (PD), and inner-outer (IO) provided by *HippUnfold.* The inner gray area corresponds to the SRLM, while the rest of the uncoloured regions correspond to the stratum pyramidale and stratum oriens layers. The white arrow in the top right inset depicts the orientation of the anterior-posterior axis. The colour of the hippocampal surface is the anterior-posterior Laplace coordinates. Finally, the black arrow depicts the intended level of the coronal slice for the cartoon depiction. (B) Simplified cartoon depiction of known microstructural circuits within the hippocampus and their approximate main orientation relative to the hippocampus, defined by the colour coded legend on the right (Zeineh et al., 2017; Nieuwenhuys et al., 2008; Duvernoy et al., 2013). (C) Primary diffusion directions for one subject (μ of the watson distribution from NODDI) overlaid on a coronal slice approximately through the hippocampal body. Coloured borders represent the hippocampal subfields provided by *HippUnfold.* (D) Pictorial example representing the NODDI and hippocampal axis vectors in a single voxel defined in (C). Cosine similarities are represented as the angle between the NODDI vector and each hippocampal vector, providing a measure of orientation coherence along each cardinal axis (see section *2.5*). Sub - Subiculum, CA - Cornu Ammonis, DG - Dentate Gyrus, SRLM - Stratum Radiatum Lacunosum Moleculare, ETC - Entorhinal Cortex.

## 2. Methods

### 2.1 Overview

A subset of 100 unrelated subjects from the publicly available Human Connectome Project (HCP) 1200 dataset were used for this study (Van Essen et al., 2013). All 100 subjects were run through *HippUnfold* (DeKraker et al., 2022), a new automated tool for surface-based subfield segmentation and hippocampal unfolding (see section *2.3*; DeKraker et al., 2018). The Laplacian coordinates generated from *HippUnfold* within each subject were used to calculate gradient vector fields along each axis of the hippocampus (see section *2.5* & Figure 1A). NODDI and DTI metrics were calculated in each subject’s native space using whole-brain diffusion images (see *2.2* and *2.4*). Cosine similarities between the NODDI orientational vector (defined as μ of the Watson distribution; see section *2.4*) and the vectors along each of the 3 axes (AP, PD, and IO) were calculated at each voxel. Furthermore, the T1w/T2w ratio was calculated as a proxy for myelin content. Macrostructural measures of curvature, gyrification, and thickness were calculated along the midthickness surface (middle of the hippocampal gray matter) of the hippocampus across all subjects. NODDI measures of ODI and NDI, DTI measures of FA and MD, and the cosine similarities were all sampled along the midthickness surface within each subject and averaged in unfolded space (DeKraker et al., 2018; DeKraker et al., 2022). Plots of NODDI and DTI metrics, cosine similarities, and macrostructure metrics across the midthickness surface were visualized as folded and unfolded surfaces. Logistic regression was performed at the level of subfield averages using the NODDI and T1w/T2w metrics to elucidate their variability across the subfields. Logistic regression was chosen due to its simplicity and interpretability. Finally, Orthogonal Projective Non-Negative Matrix Factorization (OPNNMF) was used to capture co-varying regions of the hippocampus and to examine the dimensions of macro- and microstructure hippocampal organization.

### 2.2 Data acquisition and preprocessing

We used the publicly available HCP young adult dataset (ages 22-35), which consisted of structural and diffusion MRI data for 1200 subjects (Van Essen et al., 2013). To avoid any biases caused by family structures, we chose the 100 unrelated subjects subset for analysis (mean age: 27.52 years +/- 3.47 years; F/M: 54/46). Data included T1-weighted (T1w) and T2-weighted (T2w) structural images at 0.7 mm^3^ isotropic resolution and diffusion-weighted data at 1.25 mm^3^ isotropic resolution. Structural images were obtained using a 3D MPRAGE sequence (TR-2400ms, TE–2.14ms, TI-1000ms, FOV-224×224 mm). Diffusion images were obtained using a spin-echo echo-planar imgaging sequence (b=0 (18 acquisitions), 1000, 2000, 3000s/mm^2^, 90 diffusion-encoding directions, TR-5520ms, TE-89.5ms, FOV-210×180mm). Data used in the preparation of this work were obtained from the Human Connectome Project (HCP) database (Van Essen et al., 2013). In this work we utilized the preprocessed structural and diffusion images for the HCP dataset. Preprocessing of structural images included: gradient distortion correction, coregistration and averaging of repeated T1w and T2w runs using 6-DOF rigid transformation, initial brain extractions for T1w and T2w, field map distortion correction and registration of T2w with T1w images, bias field correction, and atlas registration. Preprocessing of diffusion images included: intensity normalization across runs, EPI distortion correction, eddy current and motion correction, gradient nonlinearity correction, and registration of the mean b0 image to T1w native space. The full pre-processing pipeline for structural and diffusion images were published elsewhere (Andersson et al., 2015; Glasser et al., 2013; Jenkinson et al., 2002; Sotiropoulos et al., 2013) and can be found at the HCP website (https://www.humanconnectome.org/study/hcp-young-adult). To derive a correlate of myelin, we divided the T1w image intensity by the T2w image intensity and corrected for the bias field (Glasser & Van Essen, 2011; Glasser et al., 2014), which is referred to as T1w/T2w for the rest of the paper.

### 2.3 HippUnfold - Hippocampal unfolding and surface-based segmentation

The newly developed *HippUnfold* tool was used in the current study for surface-based segmentation and unfolding of the hippocampus (DeKraker et al., 2022). *HippUnfold* is predicated on the idea that the hippocampal subfields are topologically constrained as they differentiate from a flat cortical mantle (Duvernoy et al., 2013). Using a ‘U-net’ deep convolutional neural network (Isensee et al., 2021), *HippUnfold* provides a detailed subject-specific tissue segmentation of the hippocampal gray matter and its topological boundaries (for example, the Hippocampal Amygdala Transition Area and the Indusium Griseum as the most anterior and posterior topological boundaries, respectively) required for unfolding (DeKraker et al., 2022). Segmentation is done for each individual hippocampi preserving its topologically consistent structure, which is critical for inter-individual alignment across variably shaped hippocampi. See Dekraker et al. (2022) for more detailed information on *HippUnfold*.

The outputs of *HippUnfold* utilized in this study included the provided subfield segmentation, Laplacian coordinates for gradient field calculation (see section 2.5), macrostructure measures of curvature, gyrification, and thickness, and the midthickness surface representation for sampling volumetric space metrics onto the surface (see section 2.4). Due to the small size of the Dentate Gyrus (DG) and CA4, we combined them into a single DG/CA4 subfield label. As well, portions of the DG were excluded in our surface representation. All subfield segmentations for both hemispheres were reviewed for gross errors by BK. The midthickness surface used in this study was composed of 2004 vertices with a spacing of roughly 1mm. Averaging of each metric across subjects was performed at each vertex in unfolded space.

### 2.4 Characterization of microstructure with NODDI & DTI

NODDI models the diffusion signal as a combination from 3 microstructural environments: intra-cellular, extra-cellular, and cerebrospinal fluid (CSF) (Zhang et al., 2012). The intra-cellular compartment is considered the space that is bounded by neurites, which is modelled as a set of sticks. The stick geometry captures the restricted diffusion of water perpendicular to neurites, and the relatively unhindered diffusion along them. Furthermore, sticks can capture the wide range of neurite orientations, from highly coherent to highly dispersed tissue. The extra-cellular compartment is the space around the neurites, which consists of glial cells and in gray matter, the somas. In extra-cellular space, the signal is modeled as Gaussian anisotropic diffusion to represent the hindered but not restricted movement of water. Finally, the CSF compartment is modelled as Gaussian isotropic diffusion, representing the free movement of water. NODDI does not draw any *a priori* assumptions about whether a voxel is gray matter, white matter or CSF, and thus it treats each voxel as a possible combination of different compartments (Zhang et al., 2012). Thus, the normalized dMRI signal can be written as:

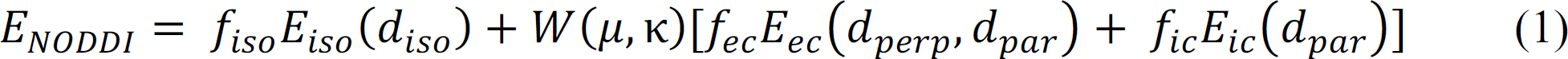

Where *f_iso_E_iso_, f_ec_E_ec_* and *f_ic_E_iC_* are the volume and signal fractions of the CSF, extra-cellular, and intra-cellular (NDI) compartments, respectively. The extra-cellular and intra-cellular compartments are linked orientationally by the Watson distribution *W*(μ, κ), where κ is the concentration parameter 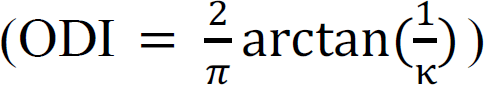 and μ is the mean orientation of the Watson distribution (herein referred to as the NODDI microstructural vector or primary diffusion direction). The hindered perpendicular diffusion of the extra-cellular compartment *d_perp_* is set via a tortuosity model. The original NODDI model which was developed mainly for white matter sets the parallel diffusivity value 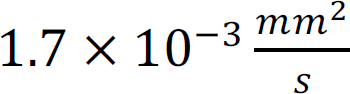 and the isotropic or CSF compartment diffusion to 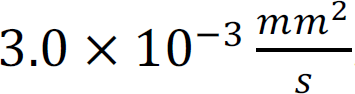. Previous studies in the gray matter have sought to optimize *d_par_* and have consistently found that the lowest mean squared error is achieved with *d_par_* equal to 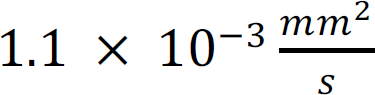 (Guerrero et al., 2019; Fukutomi et al., 2018). Thus, in the current study we used the gray matter optimized *d_par_* 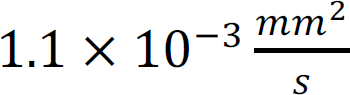 for fitting the NODDI model. The Microstructure Diffusion Toolbox (MDT; Harms et al., 2017) was utilized to fit the NODDI model using whole-brain diffusion images aligned to their respective T1w space with all b-values (b=0, 1000, 2000, 3000 s/mm^’^). The validity of the assumptions of the NODDI model are discussed in section 4.6.

We also used the MDT (Harms et al., 2017) to calculate metrics of FA and MD using DTI. DTI was performed using only the 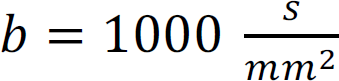 volumes to align with typical DTI experiments (Behrens & Johansen-Berg, 2014). Both the NODDI and DTI metrics were mapped onto the hippocampal midthickness surface using the process described below.

We used Connectome Workbench (https://github.com/Washington-University/workbench) to sample values at each surface vertex from volume data. In this study we used 2004 vertices defined along the midthickness surface of the hippocampus. Sampling along the midthickness surface helps reduce partial volume effects. To sample voxel data along the midthickness surface we used a ribbon-constrained mapping algorithm which also requires the inner and outer surfaces also generated by *HippUnfold.* The ribbon method constructs a polyhedron from the vertex’s neighbor on each surface defined, and then estimates the volume of the polyhedron that falls inside any nearby voxels to use as weights. We further reduced the weight of any voxel based on its distance from the midthickness surface, where the scaling value was calculated using a Gaussian with a standard deviation determined by the laminar thickness at each vertex. This had the effect of more aggressively down-weighting voxels further from the midthickness surface where the hippocampus is thinner. We then averaged each metric at each vertex across all subjects to generate the average maps which were plotted in folded and unfolded space.

### 2.5 Examining the primary direction of diffusion relative to hippocampal axes

Water molecules diffuse more readily parallel to microstructure, which in the hippocampus tends to be aligned along the AP, PD, and IO axes (Zeineh et al., 2017; Nieuwenhuys et al., 2008; Duvernoy et al., 2013). Thus, analyses were performed to examine how the primary direction of diffusion was oriented relative to these axes, with the goal of elucidating the stereotyped orientation of hippocampal microstructure (Figure 1). We obtained gradient vector fields along the AP, PD, and IO axes by taking the first derivative of the respective Laplacian coordinates provided by *HippUnfold* (Figure 1A), such that the vectors only pointed along one of the axes. That is, we computed the partial derivative of the Laplacian coordinate function along the x, y, and z spatial dimension:

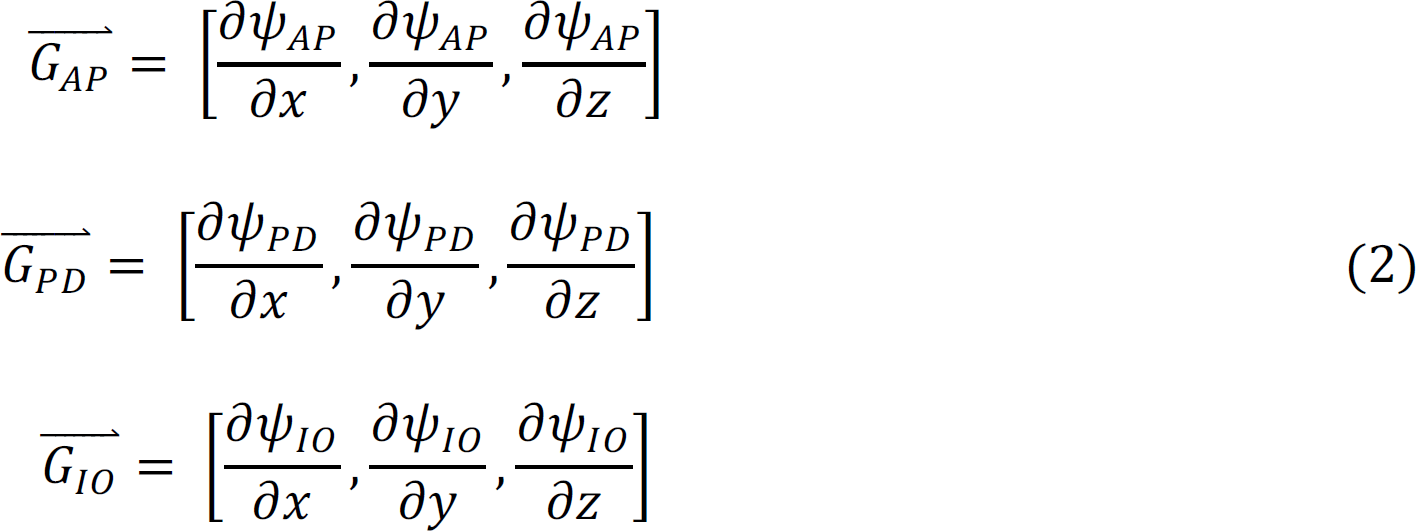

Where the function ψ_∗_ represents the spatial Laplacian coordinates along a particular hippocampal axis (AP, PD, or IO), which were calculated by solving Laplaces equation along each axis (∇^’^(ψ) = 0) (DeKraker et al., 2022). The result was 3 distinct vector images within a hemisphere for each subject and axis. With the aligned NODDI microstructural vectors (representing the primary diffusion direction), we calculated cosine similarities between the generated vectors along the AP, PD, and IO axes and the NODDI microstructural vector at each voxel (Figure 1C & D). All vectors were normalized before calculating cosine similarities. The cosine similarity was defined as the inner product between vectors:

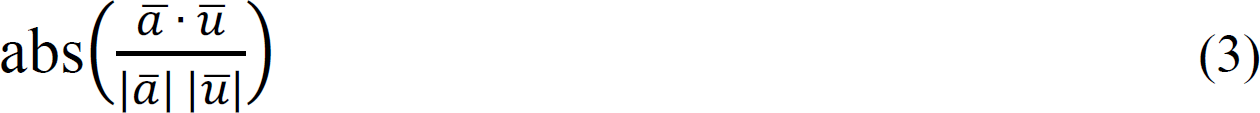

Where *ā* was a hippocampal axis vector in a single voxel (AP, PD, or IO), and *ū* was the NODDI microstructural vector (μ of the Watson distribution) at the same voxel. A higher cosine similarity meant that diffusion is increasingly oriented along that hippocampal axis (cosine similarities of 0 and 1 correspond to angles of 90 degrees and 0 degrees, respectively). The cosine similarities were then put into context of the known spatial arrangement of hippocampal microstructure and their stereotyped orientation (Figure 1A and B) under the assumption that the primary direction of diffusion modelled the main microstructural orientation at each voxel. There were a total of 3 cosine similarity images (one for AP, PD, and IO similarity values) within each hemisphere for each subject. Each scalar cosine similarity image was sampled along the midthickness surface and averaged as described above.

### 2.6 Correlations between all metrics

Spearman correlations were performed at both the subject-level and at the level of the vertex averaged maps (Figure 2C-J average maps). However, it is difficult to test the significance of the spatial correspondence between maps. One proposed solution that has been implemented using the spherical representation of the cortical surface has been to perform spin testing, where the cortical sphere is randomly permuted to derive a null distribution of association which preserves the spatial autocorrelation in the data (Alexander-Bloch et al., 2018). Critically, the geometry of our unfolded hippocampus cannot be well represented by a sphere. We thus developed a similar method on the unfolded hippocampus. Our developed method uses periodic boundary conditions on the unfolded plane, which can be thought of as performing spin testing on a torus geometry. This allows us to perform rotations and translations and thus derive permuted null distributions. We then used this hippocampal spin test to analyze the significance of the observed associations between the averaged maps. We performed 2500 permutations between any two averaged maps, and we then performed false-discovery rate correction on the derived p-values using the Benjamini-Hochberg method. Technical and practical background on the developed hippocampal spin test can be found in Supplementary Document 1, and the spin test code can be found at Karat (2023).

**Figure 2.**
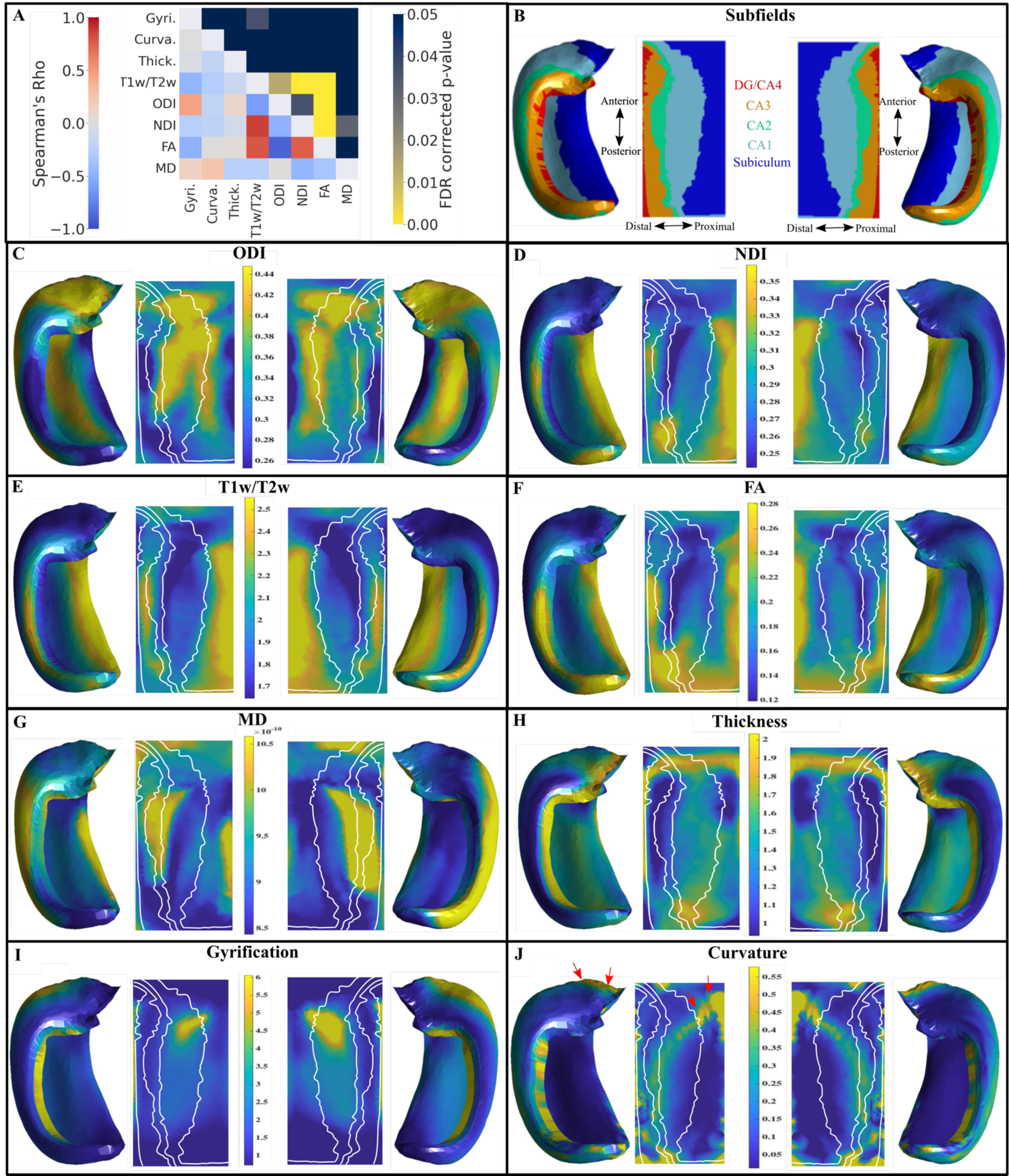
Correlations and plots of mean macro- and microstructure metrics on averaged hippocampal midthickness surfaces in folded and unfolded space for left and right hemispheres. (A) Lower triangle of the heatmap shows the Spearman’s rho correlations of the average maps after combining both left and right hemispheres. The upper triangle represents the false-discovery rate corrected p-values derived from the hippocampus spin testing using 2500 permutations. Note that the colour bar is inverted such that any brighter component of the heatmap corresponds to a significant p-value. (B) Left and right hippocampal subfields from a manual segmentation of a histological reference (Ammunts et al., 2013; DeKraker et al., 2020). Unfolded space is shown in the same orientation for left and right hemispheres. DG - Dentate Gyrus, CA - Cornu Ammonis. (C,D) Orientation Dispersion Index (ODI) and Neurite Density Index (NDI) from NODDI. White lines represent subfield borders shown in (B). (E) T1w/T2w ratio. (F,G) Diffusion Tensor Imaging metrics of Fractional Anisotropy (FA) and Mean Diffusivity (MD - *m*^2^/*s*). (H-J) Macrostructure measures of thickness, gyrification, and curvature. (J) Red arrows highlight the highly curved “spine” of the hippocampus.

### 2.7 Orthogonal Projective NNMF (OPNNMF)

Orthogonal Projective NNMF (OPNNMF) was used in this work to attempt to identify co-varying regions in the hippocampus using the metrics described above (Sotiras et al., 2015; Yang & Oja, 2010). OPNNMF decomposes an input matrix X of dimensions a x b into a component matrix C (a x k) and a weight matrix W (k x b). The number of components (k) is defined *a priori*. The component and weight matrices are derived such that their multiplication best reconstructs the input data (X ∼ C x W). OPNNMF solves the following minimization problem to estimate C (Sotiras et al., 2015):

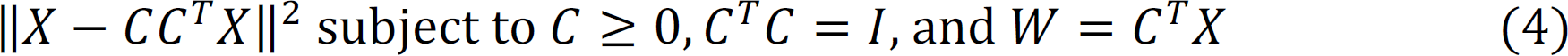

Where || ||^2^ represents the squared Frobenius norm and I denotes the identity matrix which enforces orthogonality among C. C is first initialized using a non-negative double singular value decomposition (Boutsidis & Gallopoulos, 2008). Then, C is updated through an iterative process until it converges on an optimal solution. The iterative multiplicative update rule is as reported by Yang and Oja (2010):

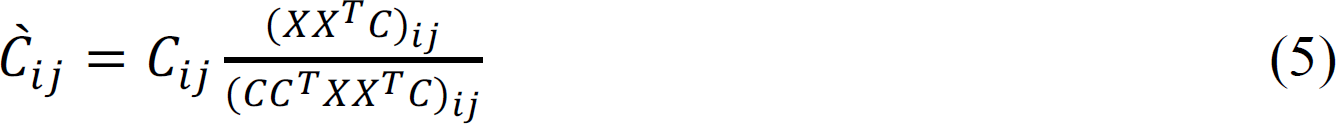

Where i represents the number of vertices and j represents the number of components. The component matrix C represents the latent structure in the data and allows for an examination of the underlying covariance in multivariate data. As done in Patel et al. (2020) the sparse and non-overlapping component matrix allows for each vertex to be assigned an output component using a winner take all method which improves the interpretability of the spatial output components. The weight matrix W represents the subject-metric coefficients, allowing for an examination of subject-specific and metric-specific contributions to each component.

### 2.8 Implementing OPNNMF

A total of 11 metrics were included in the OPNNMF implementation (ODI, NDI, T1w/T2w, FA, MD, gyrification, thickness, curvature, AP cosine similarity, PD cosine similarity, IO cosine similarity) with subsets of these metrics used for more specific analyses (i.e. NODDI only, DTI only, macrostructure only, and cosine similarity only). The input matrix X was built using all 2004 vertices of the midthickness surface in unfolded space for all 11 metrics across all 100 subjects per hemisphere. That is, each subject contributed 11 unfolded space maps to the input matrix. Thus, the input matrix had 1100 columns (100 subjects x 11 metrics - defined as subject-metrics) and 2004 rows (2004 vertices) for a single hemisphere. Normalization was required since the metrics exist on different scales. First, each metric was z-scored within each hemisphere. Then, each z-scored metric distribution was shifted by the minimum value from all the z-scored metrics to ensure all metrics were on the same scale and there were no negative values. All distributions were manually inspected to ensure the minimum value used was not an outlier.

OPNNMF was implemented using publicly available and open MATLAB code at https://github.com/asotiras/brainparts (Sotiras et al., 2015; Yang & Oja, 2010; Boutsidis & Gallopoulos, 2008; Halko et al., 2011). OPNNMF was run with a max number of iterations = 10000, tolerance = 0.00001, and non-negative double singular value decomposition initialization.

### 2.9 Stability & Reconstruction Error

The quantification of OPNNMF decomposition stability followed that of Patel et al. (2020). Stability was assessed by examining the similarity of the spatial component matrix C across varying splits of data. All 100 subjects were randomly split into two groups of equal sizes. OPNNMF was then performed on each split independently. A within-split similarity matrix was then derived by multiplying a particular splits component matrix by the transpose of itself (i.e. a cosine similarity). The result is a 2004×2004 (number of vertices x number of vertices) matrix where each row contains the cosine similarity of component scores between a vertex and all other vertices. Finally, a Pearson’s correlation coefficient was calculated across the rows of the cosine similarity matrix between splits to quantify if the decomposition maintained the relationships between vertices. The above process was repeated 6 times, each with a new random split of the data. The mean and standard deviation of the correlation coefficient was taken across all vertices and splits for a given component solution. A correlation of 1 represented perfect stability (i.e. each split of the data had perfect correspondence between vertex relationships), whereas -1 represented instability. The above process was then repeated for different component decompositions, from k=2 to k=12. Reconstruction error was calculated through 3-steps. First, the component matrix C and the weight matrix W were estimated and then multiplied together to return the reconstructed input matrix. The original and reconstructed input matrix were then subtracted to obtain a reconstruction error matrix. The Frobenius norm of the reconstruction error matrix was then taken to get the reconstruction error. The gradient in the reconstruction error was taken across solutions with varying component numbers to assess the magnitude of the improvement in reconstruction error when adding more components.

### 2.10 Interpreting OPNNMF

The output component matrix C contains a component value for each vertex while the weight matrix W describes how each subject-metric is projected onto each component. A large value in the component matrix can be interpreted as a particular vertex being identified as a part of the variance pattern. The weight matrix can be used to elucidate which metrics contributed to each component as well as inter-subject variance within metrics. In the current study these 2 matrices were used to explore spatial patterns and the contributions of particular metrics to each component. A winner-take-all method was used where a vertex was assigned the integer of the component with the highest component weighting value from the matrix C. The matrix W was z-scored within each component row and then plotted to examine metric-specific trends.

## 3. Results

The results begin with qualitative descriptions of average macro- and microstructural measures and their correlations on the midthickness hippocampal surface (middle of the hippocampal gray matter). We also show the significance of these correlations using the developed hippocampal spin test. We then present results from a logistic regression model used to examine the separability of the subfields using microstructural features. We then show the cosine similarities between NODDI microstructural vectors and the hippocampal axis vectors. Finally, we present the OPNNMF results including a stability analysis and a 6-component solution.

### 3.1 Distributions of Hippocampal Metrics

Figure 2 presents Spearman’s rho correlations, mean macro- and microstructural metrics, along with subfield segmentations shown on an averaged hippocampal midthickness surface in folded and unfolded space. The standard deviation of these metrics is shown in supplementary Figure 1. The orientation dispersion (ODI - Figure 2C) is highest in the anterior and body of CA1 and the distal parts of the subiculum, while dispersion is lowest in the body and posterior of DG/CA4, CA3 and CA2 and at the most proximal edge of the subiculum. Neurite density (NDI - Figure 2D) is highest in the body and tail of the subiculum, while there is lower neurite density in CA1 and the DG/CA4. The T1w/T2w ratio has a strikingly similar distribution to that of NDI, as found in previous cortical studies (Figure 2E; Fukutomi et al., 2018). T1w/T2w, ODI, and NDI maps appear to vary across the subfields.

Macrostructure features of thickness, gyrification, and curvature are shown in Figure 2H-J. Gyrification is largest in anterior CA1 and the DG/CA4. The thickest regions are the anterior and posterior of the subiculum and CA1, as well as throughout the DG/CA4, while CA3 and CA2 are thin. Curvature tends to be highest in the anterior part of the subiculum, along the spine of the hippocampus (red arrows in Figure 2J), and in CA3. These findings are largely in line with previous work (DeKraker et al., 2020).

Spearman’s rho correlations between averaged maps of these metrics can be seen in the lower triangle of Figure 2A while the false-discovery rate (FDR) corrected p-values derived from the hippocampal spin test can be seen in the upper triangle. Similar correlations between all maps at the subject level can be seen in supplementary Figure 2. The null distributions given by the spin test using the averaged maps can be seen in supplementary Figure 3.

### 3.2 Correlations between NODDI metrics & T1w/T2w

In Figure 2D and E, strong qualitative similarities can be seen between NDI and T1w/T2w. Both are high in the subiculum and CA3/CA2 regions, while hypointensities are noted in CA1. A significant spatial overlap is seen between the positively correlated NDI and T1w/T2w maps (ρ = 0.86, FDR corrected p-value < 0.05). ODI (Figure 2C) and T1w/T2w (Figure 2D) are also significantly overlapping and are negatively correlated (ρ = -0.61, FDR corrected p-value < 0.05). A similar correlation has also been noted across the entire cortex (Fukutomi et al., 2018).

### 3.3 Correlations between NODDI & DTI metrics

Qualitatively, the map of FA (Figure 2F) resembles the NDI map (Figure 2D) and the inverse of the ODI map (Figure 2C). Particularly, the distinct pattern of high dispersion equates to low FA, while high neurite density equates to high FA. Furthermore, the region of low neurite density (Figure 2D) in the body of CA1 and CA2 corresponds to a region of high MD (Figure 2G). ODI and FA have a significant spatial overlap and are negatively correlated (ρ = -0.88, FDR corrected p-value < 0.05). Furthermore, NDI and FA are significantly overlapping and are positively correlated (ρ = 0.75, FDR corrected p-value < 0.05) and NDI and MD are significantly overlapping and negatively correlated (ρ = -0.46, FDR corrected p-value < 0.05).

A disentangling of FA as determined by ODI and NDI has been reported previously in cortical gray matter (Zhang et al., 2012). Two voxels with different neurite densities can have the same FA if the one with the larger neurite density also has the larger dispersion (Zhang et al., 2012). That is, anisotropy as measured with FA conflates neurite density and orientation dispersion. We report a similar level of disentangling within the hippocampus in supplementary Figure 4, where within any bin of FA there is a range of potential ODI and NDI values.

### 3.4 Correlations between macrostructure and NODDI metrics

The only significant overlap between a macrostructural and microstructural map is gyrification and T1w/T2w, which are negatively correlated (ρ = -0.46, FDR corrected p-value < 0.05). Regions of high gyrification in CA1 spatially correspond to regions of low T1w/T2w content. While there are some other notable correlations, such as a moderate positive correlation between gyrification and ODI (ρ = 0.45) and a moderate negative correlation between gyrification and FA (ρ = -0.49), none were significantly overlapping. Correlations between NODDI and macrostructural measures at the subfield-averaged level can be seen in supplementary Figure 5.

### 3.5 Evaluating the variability of the microstructural metrics across the subfields

To examine if the microstructural metrics used here could provide differentiability to the subfields, we trained a simple logistic regression classifier with L2 regularization on subfield averaged microstructural measures using the scikit-learn (version 1.1.2) package in python (Pedregosa et al., 2011). 70 subjects were included in the training set, while 30 subjects were included in the testing set (both hemispheres were combined). ODI, NDI, and T1w/T2w averages across the midthickness surface within each of the 5 subfields per hemisphere were obtained within each subject in both the training and testing set. A logistic regression model was then trained to predict the subfield label given the subfield averaged microstructural metrics. This model was then tested on the unseen data from the testing set to quantify if the microstructural metrics are differentiable across the subfields. Examining all the data points for ODI, NDI, and T1w/T2w, colour coded by subfield class, it appears that the subfields present as relatively separable clusters apart from CA2 which appears to have lots of microstructural overlap with the other subfields (Figure 3A). The same clustering can be seen in the test set (Figure 3B) left out to evaluate the performance of the logistic regression classifier (Figure 3C). Using a confusion matrix and F1-scores defined as (2*precision*recall) / (precision+recall), it can be seen that the simple logistic regression classifier was able to perform well in delineating the subiculum (F1-score = 0.87), CA1 (F1-score = 0.85), CA3 (F1-score = 0.69), and the DG/CA4 (F1-score = 0.83) using microstructural measures of ODI, NDI, and T1w/T2w (Figure 3D). However, CA2 was not as differentiable from the rest of the subfields using these metrics, with an F1-score of 0.53. Furthermore, we found that ODI and NDI generally had higher F1-scores and greater separability across the subfields then the DTI metrics of FA and MD (supplementary Figure 6).

**Figure 3.**
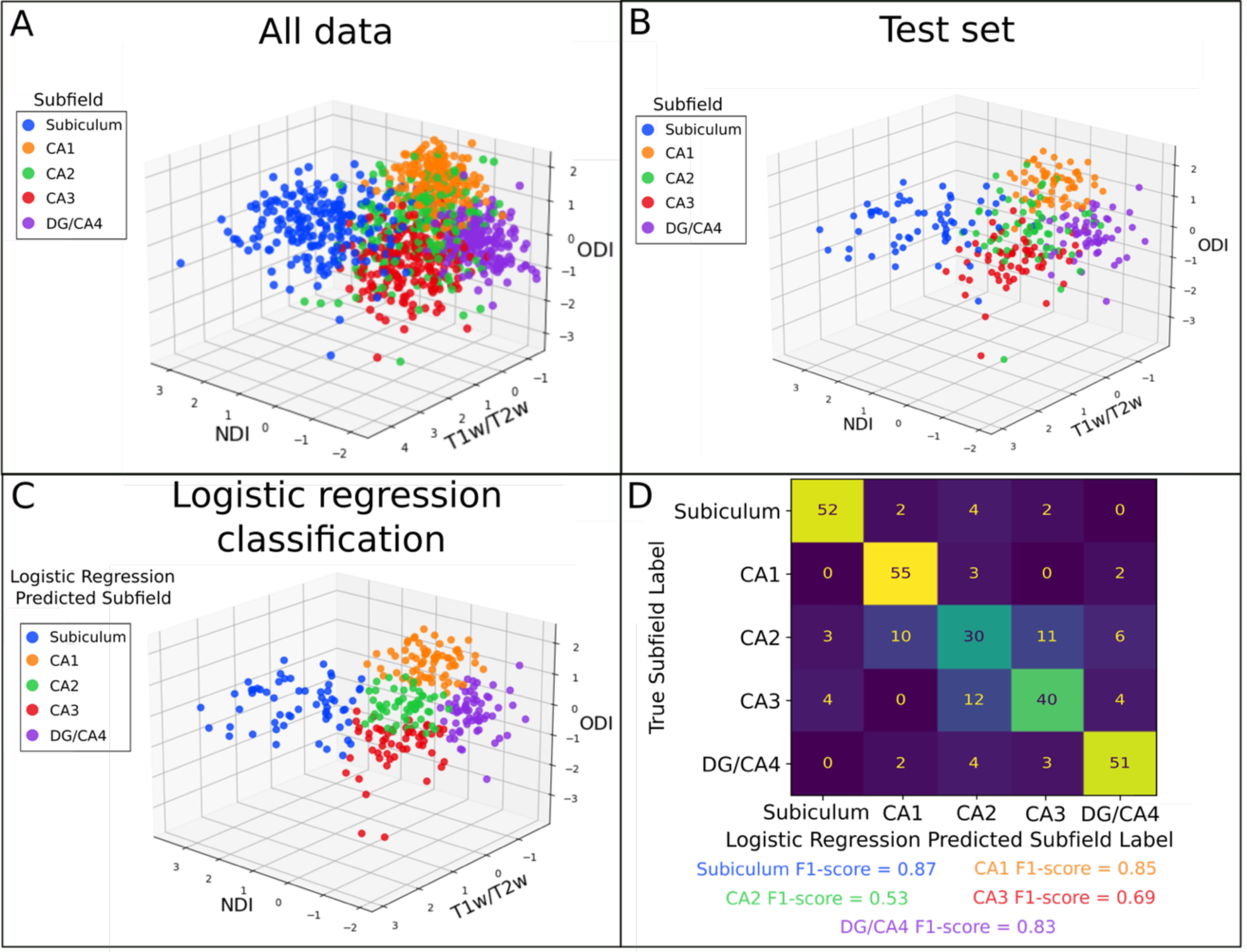
The performance of a logistic regression classifier to capture microstructural variability across the subfields. (A) Subfield averaged ODI, NDI, and T1w/T2w from all subjects and hemispheres combined coloured by subfield label. (B) Test set data from 30 subjects coloured by subfield. (C) Logistic regression classified labels on the test set seen in (B). (D) Confusion matrix and subfield-specific F1-scores to evaluate logistic regression performance on classifying the subfields using the microstructural measures of the test set.

### 3.6 Examination of the primary direction of diffusion relative to hippocampal axes

This section qualitatively analyzes the mean of the cosine similarities (Figure 1D; Figure 4) between the hippocampal vectors along the anterior-posterior (AP), proximal-distal (PD), and inner-outer (IO) (Figure 1A) axes and the NODDI microstructural vectors along the midthickness surface (Figure 1C).

**Figure 4.**
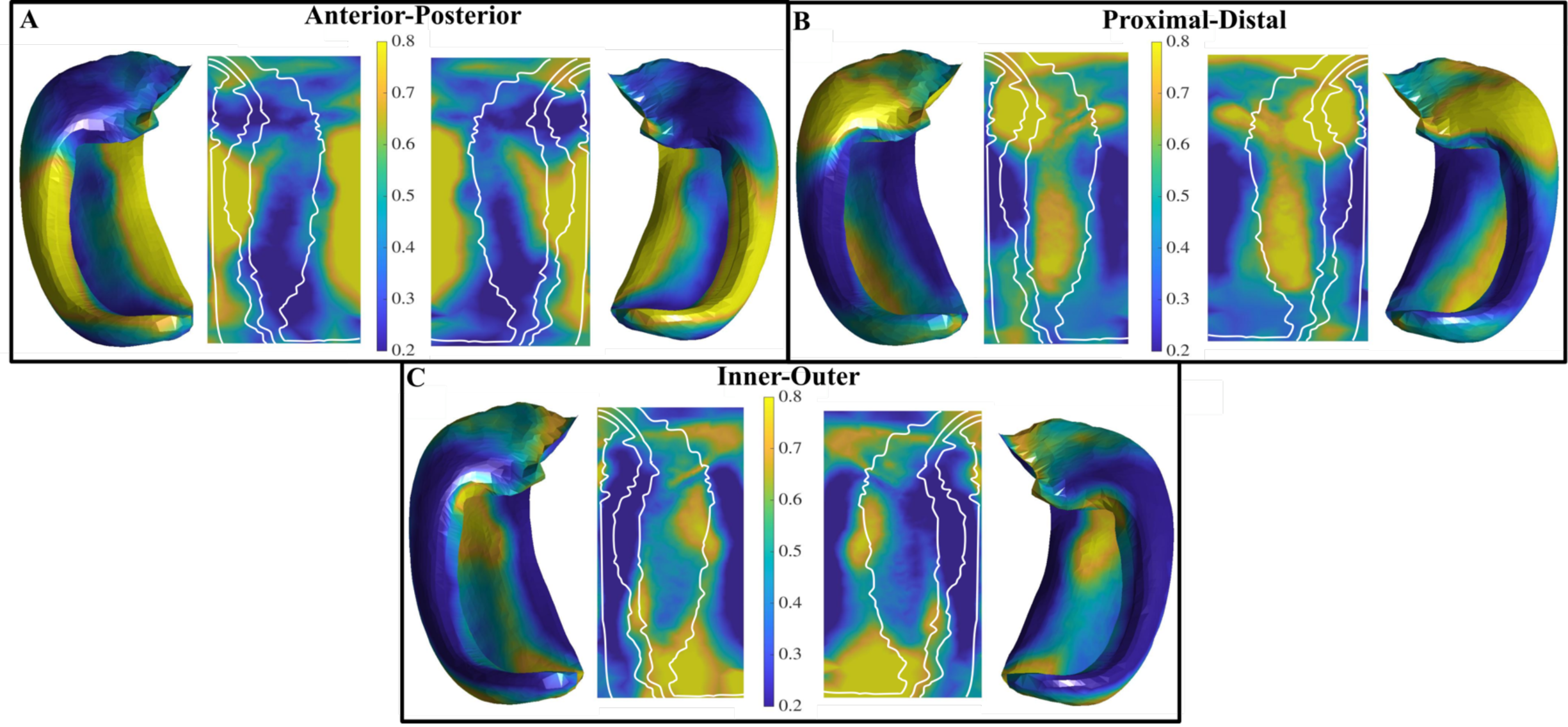
Mean of the cosine similarities between hippocampal axis vectors and NODDI microstructural vectors in the left and right hemisphere along the midthickness surface. High cosine similarities correspond to a high alignment of the NODDI microstructural vector along that particular hippocampal axis. (A) Distribution of cosine similarities along the anterior-posterior direction. (B) Distribution of cosine similarities along the proximal-distal (tangential) direction. (C) Distribution of cosine similarities along the inner-outer (laminar) direction.

High AP alignment can be seen in the body of the DG to CA3, as well as in the subiculum. Furthermore, there is relatively little AP oriented diffusion in CA1.

High PD oriented diffusion can be seen in the head of CA3, CA2, CA1, and in the body of CA1. There was little PD oriented diffusion in the body of DG/CA4, CA3, CA2, and the subiculum.

High IO alignment can be seen in the tail of the subiculum and CA1, as well as throughout the body of CA1, while low IO oriented diffusion can be seen in the body of the DG/CA4, CA3, CA2, and the subiculum.

The cosine similarities varied the greatest in CA1 and the subiculum across AP, PD, and IO directions (supplementary Figure 7). Noticeable differences in the orientation of diffusion across the subfields can be seen in supplementary Figure 8.

### 3.7 Stability Analysis

The results of the stability analysis can be seen in Figure 5. Figure 5A presents the stability and the gradient in the reconstruction error using all the metrics combined that are shown in Figure 5B-E. The goal of the stability analysis was to elucidate the largest component value that was still stable and provided a relative gain in reconstruction error. In Figure 5A it can be seen that a component solution of k=6 has good stability with relatively low standard deviation.

**Figure 5.**
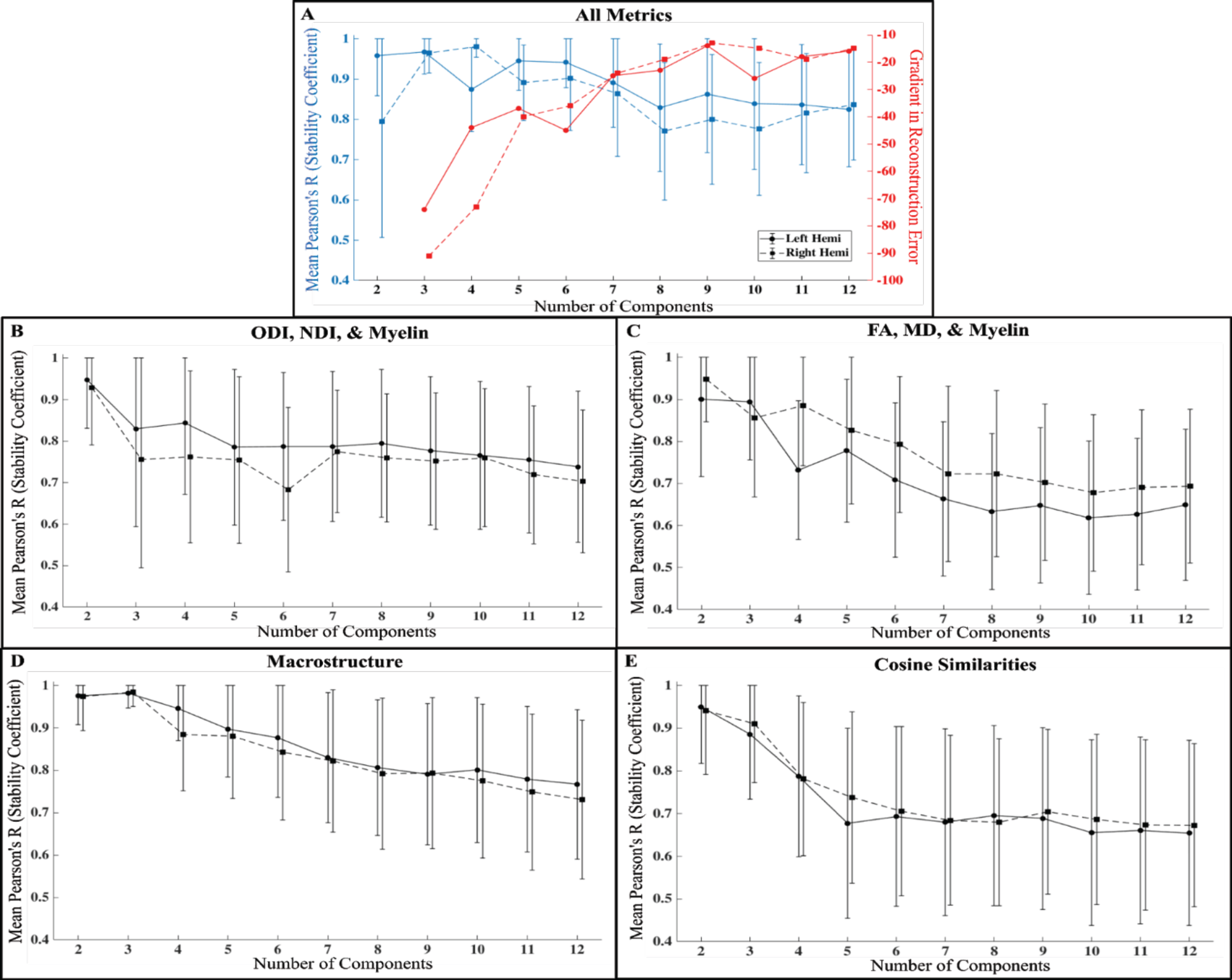
Stability coefficient and the gradient in reconstruction error based on the number of components used for the OPNNMF solution. Filled in circles plus solid lines are the left hemisphere and filled in squares plus dotted lines are the right hemisphere. Error lines show +/- 1 SD. (A) Stability coefficient (blue) and the gradient in reconstruction error (red) as a function of the number of components using all metrics for NMF in B-E. (B) Stability coefficient for NODDI metrics (ODI and NDI) plus T1w/T2w. (C) Stability coefficient for DTI metrics (FA and MD) plus T1w/T2w. (D) Stability coefficient for macrostructure metrics (gyrification, thickness, curvature). (E) Stability coefficient for cosine similarities (AP, PD, and IO). Points between hemispheres are slightly offset along the x-axis so that error bars are visible.

Comparatively, decomposing into a larger number of components decreases the stability of the OPNNMF solution. As well, k=6 does provide a relative gain in reconstruction error, although the largest gain in reconstruction error occurs when moving from a k=2 to a k=3 solution. The stability analysis suggests that k=6 is the highest component value that is largely stable, thus we use this for the decomposition results using all metrics.

Another goal of the stability analysis was to compare the stability of the decomposition using all metrics versus using smaller groupings of metrics, such as NODDI (ODI and NDI) plus T1w/T2w. Comparing Figure 5A with Figure 5B-E, it can be seen that for almost all component values the all metric solution tends to be more stable than any of the smaller metric groupings. This is especially true for the larger component values above k=6. These results suggest that the use of multiple metrics results in more stable parcellations, as found in Patel et al. (2020).

The 6-component solution using all metrics is presented in Figure 6 for both the left and right hippocampus. 4, 5, 6, and 7-component solutions using all the metrics can be found in supplementary Figure 9. 4-component solutions for all smaller metric combinations shown in Figure 5B-E can be found in supplementary Figure 10.

**Figure 6.**
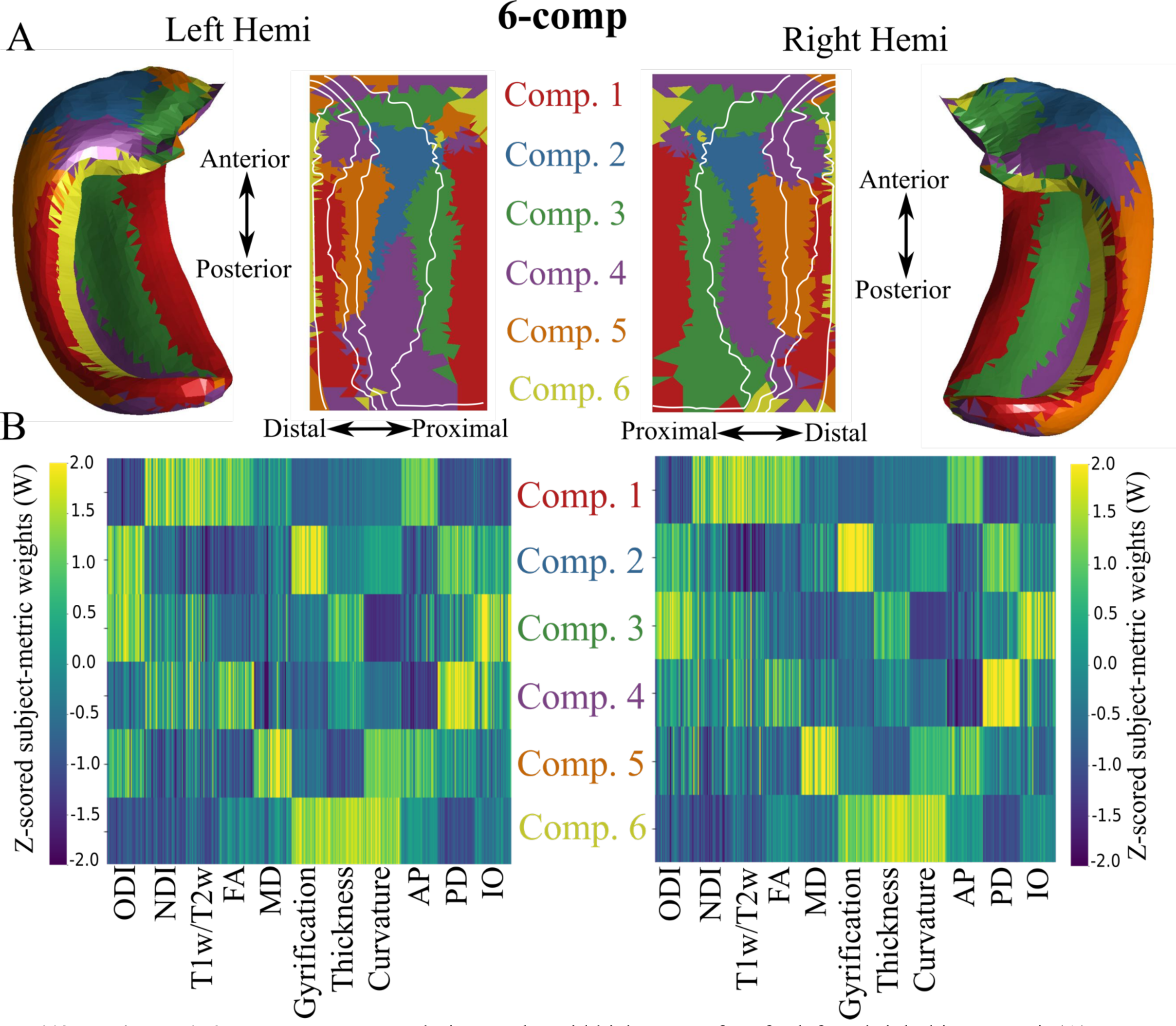
6-component NMF solution on the midthickness surface for left and right hippocampi. (A) Winner-take all output at each vertex shown in folded and unfolded space. White lines denote subfield borders. (B) Z-scored subject-metric weight matrices across each of the 6 components, denoting the z-scored contribution of each metric to each component. AP, PD, and IO represent the 3 cosine similarity metrics.

### 3.8 Description of the 6-component Solution

Figure 6A depicts the winner-take-all method applied at each vertex in folded and unfolded space for 6-components. Figure 6B shows the z-scored subject-metric weight matrices. In the following paragraph we describe the first 3 components including their location relative to the subfields (proximal-distal/medial-lateral axes) as well as along the anterior-posterior (longitudinal) axis. We then briefly describe components 4-6. We also describe the features which contribute to each component. The left and right hippocampus do have similar covariance patterns although the component ordering differs. For example, the vertices of left component 1 correspond to the vertices of right component 2. To simplify our descriptions, we take the ordering of the left hippocampus for the rest of the paper, and we have made this adjustment such that the difference in ordering does not need to be considered for Figure 6 (Patel et al., 2020).

Component 1 is characterized by a cluster of vertices through the body and tail of the most proximal edge of the subiculum and through the body and tail of CA3. This component spans around the bottom two-thirds of the hippocampus across its anterior-posterior axis. Component 1 is characterized by high NDI, T1w/T2w, FA, and AP cosine similarity, with lower ODI, PD and IO cosine similarity. This may reflect the large anisotropic AP oriented fiber bundles that are myelinated such as the cingulum bundle for the proximal edge of the subiculum and the fimbria for CA3.

Component 2 is characterized by vertices that are present only in the body or middle one-third along the anterior-posterior axis of CA1. This component is characterized by high ODI, gyrification, and PD and IO cosine similarities. This likely reflects a high heterogeneity in fiber orientation in CA1.

Component 3 is characterized by vertices that cross the subiculum and CA1 in a proximal-distal fashion in the head of the hippocampus, as well as vertices that span the anterior-posterior body of the hippocampus at the border between the subiculum and CA1. This component is characterized by high ODI, thickness, and IO cosine similarity.

Component 4 corresponds to a large posterior cluster that stretches into the body of CA1 characterized by a moderate range of metrics with the largest being the PD cosine similarity and FA. Component 5 corresponds to a cluster of vertices in the body of CA2 and CA1 and is represented by a large MD, curvature, and AP cosine similarity and low NDI, T1w/T2w, FA, and thickness. Interestingly, component 6 is characterized by a thin cluster of vertices roughly corresponding to the anterior and body of the DG/CA4 region. This was represented by high macrostructural measures of thickness, gyrification, and curvature and low weights for all other metrics.

A general pattern is noticed when examining the whole 6-components rather than looking at its parts. Generally, more parcellations exist along the proximal-distal direction than do in the anterior-posterior direction.

## 4. Discussion

In the current study we examined the microstructure of the hippocampus using the in vivo HCP dMRI and structural data, along with a novel surface-based method for subfield segmentation. We found that ODI was highest in the CA1 subfield, likely capturing the large heterogeneity of tangential and radial fibers. NDI and T1w/T2w were found to be strongly correlated and were highest in the subiculum and lowest in CA1 and the DG/CA4, suggesting that NODDI is likely sensitive to the myelin content of the hippocampus. Using these microstructural measures, we found that the cytoarchitectonic defined subfields were largely separable using a simple logistic regression model. OPNNMF components appeared to capture unique co-varying clusters within the hippocampus, with high medial-lateral variability. Finally, we showed distinct regions of similar microstructural orientations by examining the main direction of diffusion relative to the three hippocampal axes, which may correspond to specific microstructural properties.

### 4.1 Dispersion of neurites in the hippocampus may reflect heterogeneous radial and tangential neurite components

The Orientation Dispersion Index (ODI) is meant to characterize the variation in diffusion orientations around a single dominant direction at every voxel. A previous study using ODI and patch-wise circular variance measured using histology (measures variability in neurite orientations) has shown that both measures have lower dispersion in demyelinated lesions in patients with multiple sclerosis, where there is reduced geometrical complexity of neurites (Grussu et al., 2017). The hippocampal gray matter has a general distribution of microstructure that is similar to the neocortex, with tangential (proximal-distal) and radial (inner-outer) components that follow the curvature of the hippocampus. In the current study we showed that CA1 had the largest ODI, suggesting that it had the largest heterogeneity in neurite orientations. CA1 has large tangential neural processes, like the Schaffer collaterals and perforant path, as well as a large (yet dispersed) radial pyramidal cell layer (Duvernoy et al., 2013). By measuring the orientation of the main direction of diffusion relative to the three hippocampal axes in CA1 (Figure 4), we found either high tangential or radial diffusion, supporting the idea that ODI reflected the heterogeneity of these components. Conversely, ODI was lower in DG/CA4, CA3, and at the most proximal edge of the subiculum. In these regions the primary diffusion direction was minimally tangential or radial, and was mainly anterior-posterior or oblique with large macroscopic diffusion anisotropy (i.e. large NDI and FA). The apparent reduction in orientational heterogeneity (less radial/tangential components) and a resulting increase in anisotropy may potentially explain the low ODI in these regions. In the DG/CA4 and CA3 region this could be a result of partial voluming with the nearby fimbria, and in the subiculum it could be due to partial voluming with the nearby cingulum bundle or the perforant path at its most proximal edge (supplementary Figure 11). As hypothesized in the rest of the cortex (Fukutomi et al., 2018), it is likely that ODI in the hippocampal gray matter is largely driven by the heterogeneity of radial and tangential neurite components.

### 4.2 Hippocampal neurite density is highly correlated with T1w/T2w

The distribution of the NDI and T1w/T2w across hippocampal gray matter was similar, as seen in Figure 2D and E and as shown by their strong positive correlation and significant overlap. While the diffusion signal is generally agnostic to water within myelin, previous work has shown that myelinated axons restrict diffusion to a greater degree than unmyelinated axons (Behrens & Johansen-Berg, 2014). This increase in restriction due to myelin would result in an increase in NDI since there is more “stick” like diffusion occurring (i.e., a monoexponentially decaying signal with a slope defined by the parallel diffusivity). Furthermore, due to the short exchange time of water within dendrites and glia to the extra-cellular space, it is likely that any restricted diffusion would be reflective of myelinated axons (Jespersen et al., 2010). This suggests that NDI reflected the myelin content of the hippocampus. The T1w/T2w content and NDI were largest in the body and tail of the subiculum. High myelin content in the subiculum has been noted previously with histology (Ding & Van Hoesen, 2015). Furthermore, it is likely that the white matter of the cingulum bundle or perforant path contribute to the large T1w/T2w content seen in the subiculum. Conversely, T1w/T2w and NDI were lower in CA1, which is likely a result of a relatively sparse layer of pyramidal cells along the midthickness surface or the largely unmyelinated Schaffer collaterals (Jürgen et al., 2011; Szirmai et al., 2012). Overall, the distribution of T1w/T2w found here agreed with previous studies (DeKraker et al., 2018; Ábrahám et al., 2012). A strong positive correlation between NDI and T1w/T2w was found previously across the cortex. However, the hippocampus was found to have high values in NDI but low values of T1w/T2w when compared to the rest of the cortical areas (Fukutomi et al., 2018). Here we showed that a strong correlation between NDI and T1w/T2w still exists in the hippocampus when comparing them at a finer spatial scale. This correlation is further corroborated by another cortical study at high ex vivo resolutions in the rodent brain, in which cortical NDI was strongly correlated with staining intensity of myelinated axons (Jespersen et al., 2010). Histological work has found similar correlations in white matter, where myelin content was found to be strongly correlated with axon count (Schmierer et al., 2007). However, a recent study utilizing a multicomponent relaxometry method for imaging myelin water fraction found no significant correlation between myelin and NDI measured using NODDI in most white matter structures (Qian et al., 2020). While NDI and T1w/T2w as a proxy of myelin do appear to be correlated in gray matter including the hippocampus, further work is needed to examine this correlation in other white matter structures, including white matter surrounding the hippocampus such as the fimbria, fornix, and alveus.

### 4.3 Microstructure metrics systematically vary across the subfields

Microstructural metrics such as intra-cortical myelin and macrostructural cortical thickness have been shown to be useful in parcellating the neocortex into subregions (Nieuwenhuys, 2013; Glasser et al., 2014; Glasser & Van Essen, 2011). Furthermore, using non-negative matrix factorization of T1w/T2w, MD, and FA it was found that a 4-component solution qualitatively resembled hippocampal subfield borders (Patel et al., 2020), suggesting that T1w/T2w and microstructure may provide sufficient separability to parcellate hippocampal subfields. In the current study, by examining their averaged maps, we found that T1w/T2w, NDI, and ODI appeared to be separable qualitatively across the subfields (Figure 2). Quantitatively, we showed that a simple logistic regression model performed well in predicting the subfield label based on the subfield-averaged metrics of T1w/T2w, NDI, and ODI. This result suggests that these microstructural measures are potentially sensitive to the known microstructural differences across the subfields as defined via cytoarchitectonics. Critically, it appeared that NDI and ODI provided more separability across the subfields then FA and MD, suggesting that NODDI may be more useful than DTI in capturing known microstructural differences across subfields (supplementary Figure 6). CA2 appeared to be the exception, with a distinct pattern of MD when compared to the other subfields, while this separability of CA2 was not captured with NODDI metrics (supplementary Figure 6). Myelin content has been demonstrated previously to closely correspond to averaged subfield borders (DeKraker et al., 2018). To a lesser extent, macrostructure appeared to also follow the subfield borders, which has been noted previously for thickness (DeKraker et al., 2018). While thickness is consistently low in CA3 and CA2, and gyrification is consistently high in CA1, these measures alone may not differentiate all subfield boundaries. Thus, a combination of NODDI and macrostructural measures may provide complimentary information needed for subject-specific subfield delineation.

### 4.4 The orientation of diffusion relative to the hippocampus may be useful in identifying hippocampal microstructure

We quantified the main direction of diffusion relative to the 3 main hippocampal axes which microstructure tends to align closely with. Here we provide descriptions of microstructure (Figure 1B) that likely contribute to the orientation results. The following three paragraphs describe the anterior-posterior (AP), proximal-distal (PD), and inner-outer (IO) alignment, respectively.

The high AP alignment in the body of the DG to CA3 was likely driven by the neighbouring fimbria, the largest bundle in that region oriented AP. High AP alignment in the subiculum was likely caused by the cingulum, a large fiber bundle that traverses the parahippocampal gyrus. Some partial voluming from the outer (where the cingulum exists) to the midthickness surface was expected, which may drive this alignment.

High PD alignment in the head of CA3 was expected to be either Schaffer collaterals which curve immediately PD off of the apical dendrites of the pyramidal cells or from perforant projections coming from the entorhinal cortex and entering CA3 which are also oriented PD. High PD alignment in CA1 was likely a result of the Schaffer collaterals. The Schaffer collaterals make synaptic contact at the apical and basal dendrites of CA1 in a PD fashion (Nieuwenhuys et al., 2008; Swanson et al., 1978). However, the perforant path could have also contributed to a higher PD alignment as it moves from the entorhinal cortex to the DG synapsing on CA1 along the way (Nieuwenhuys et al., 2008). Furthermore, a high PD cosine similarity in the CA3 region could have been a result of partial voluming with the alveus, a highly PD oriented bundle that sits atop the hippocampal gray matter.

High IO alignment seen in CA1 was likely a result of the pyramidal neurons. The pyramidal somas exist in the stratum pyramidale layer of the midthickness surface, and are generally scattered in CA1 (Nieuwenhuys et al., 2008). Their axons and basal dendrites move IO towards the alveus/outer surface and their large apical dendrites move IO towards the stratum radiatum/inner surface. All IO alignment seen in CA1 was expected to be caused by the pyramidal neurons or other afferent CA1 paths such as the Schaffer collaterals which may curve IO before making contact with the apical dendrites of the pyramidal neurons. High IO alignment in the subiculum was also likely caused by pyramidal neurons as in CA1. The cosine similarities across subjects varied the greatest in CA1 and the subiculum across AP, PD, and IO directions (supplementary Figure 7).

Typical hippocampal microstructural analyses average scalar diffusion metrics (such as FA, MD, NDI, etc.) either across whole hippocampi (van Uden et al., 2015; Salmenpera et al., 2006) or whole subfields (Radhakrishnan et al., 2020), which are inherently non-specific towards microstructure that exists within and across subfields and the hippocampal long-axis. However, tractography analyses which aim to capture the continuous intra-hippocampal circuitry are difficult to perform, as at lower resolutions, tracts can be spurious requiring complex acquisition and correction schemes (Zeineh et al., 2012). The orientational analyses described here have the potential to increase specificity at in vivo resolutions by leveraging the known anatomical orientations of hippocampal microstructure. Furthermore, capturing the essence of hippocampal microstructural orientations with vertex-wise scalar values can make qualitative observations and statistical analyses more tenable than with the complex 3D orientations provided by tractography. Future studies could relate the hippocampal axis vectors to other dedicated methods of diffusion orientation representation that have the ability to faithfully represent more complex fiber configurations.

Applications of the proposed orientational methods may be useful to identify microstructure deterioration in disease states, where affected microstructure may be less prominent, and may appear as smaller cosine similarities along a particular axis. For example, perforant path lesions in rats caused rapid memory loss which was akin to early-stage Alzheimer’s disease (Kirkby & Higgins, 2001). A 2010 study found deterioration of the perforant path in aged humans using diffusion tensor imaging (Yassa et al., 2010). Perforant path degradation should result in less attenuation of the diffusion signal along its length, which may potentially show up as smaller PD cosine similarities specifically in the subiculum, CA3, and CA1, as there should be less PD oriented diffusion. This may be possible for other neurological diseases where specific microstructure is affected, such as pyramidal neuron degradation which should result in smaller IO cosine similarities. However, to draw such conclusions, further ex vivo validation with the ability to measure more ground-truth microstructural orientations will be essential to evaluate the usefulness of this method. As well, future studies will have to evaluate the efficacy of this method at clinical resolutions.

### 4.5 6-component OPNNMF solution displays distinct co-varying regions of macro- and microstructure

The hippocampus is believed to have two main interacting dimensions of organization along its medial-lateral/proximal-distal or subfield axis, and across its long or anterior-posterior axis (see Genon et al., 2021 for review). In the current study with a 6-component OPNNMF solution we found a gradient in the clusters along the proximal-distal direction, suggesting that regions along this axis had disparate macro- and microstructural properties. That is, the macro- and microstructural “axes” of the hippocampus as defined by the metrics used here tended to vary along the same axis that the subfields were defined. Variability along this axis was expected, as the subfields show differences in morphology, cytoarchitectonic profiles (Duvernoy et al., 2013; Ding & Van Hoesen, 2015), and connectivity (Andersen et al., 1971), which likely manifest as changes in macro- and microstructure. However, it is notable that a wide array of disparate metrics at in vivo resolutions show such patterns. Recently there has been interest in the long-axis organization of the hippocampus, with evidence coming from anatomical and physiological recordings in rodents (Chase et al., 2015). In the current study there appeared to be a smaller gradient in the clusters along the anterior-posterior axis, suggesting that the metrics used here varied less along the long-axis when compared to the proximal-distal axis. While functional studies have found stark variability across the long axis, it is unclear to the extent which the hippocampal microstructure varies across this axis (Genon et al., 2021, Plachti et al., 2019). Future work could look to examine the variability of dMRI measures across the long axis at higher resolutions.

### 4.6 Limitations

There are some limitations of the current research that largely pertain to the metrics used here. As noted in the introduction, DTI aims to capture the macroscopic diffusion anisotropy at each voxel, assuming that the diffusion process can be well characterized by a Gaussian distribution. Due to its popularity in both clinical and research settings, we sought to examine the distribution of two common DTI measures (FA and MD) in the hippocampus. However, in most gray matter regions there tends to be a complex arrangement of fiber orientations that cannot be well modelled via a single tensor. This can result in drastically understated FA values with generally spherical or planar DTI ellipsoids which contain minimal information on diffusion orientations (Campbell et al., 2005). While DTI has been used extensively in the hippocampus, it has generally been purported to be sensitive but not specific to its microstructure (Coras et al., 2014). In ex vivo tissue at microscopic resolutions DTI metrics of FA and MD have shown good contrast to the hippocampal laminae, demonstrating their sensitivity (Coras et al., 2014; Shepherd et al., 2007; Stolp et al., 2018). However, at lower in vivo resolutions used here a plethora of microstructure partial voluming within each voxel is inevitable, which leads to a diffusion signal that may show little macroscopic diffusion anisotropy and thus cannot be captured by a single tensor. Indeed, here we found that even the largest FA values of the hippocampus in regions of purportedly high T1w/T2w tended to range around 0.28, a relatively low amount of macroscopic anisotropy. Furthermore, it is unclear the extent to which extra-hippocampal white matter such as the fimbria and angular bundle might influence our measures sampled along the middle of the hippocampal gray matter.

The NODDI model has also been used extensively in both healthy and diseased states to provide metrics that are biologically grounded. Such work involved explicitly modelling the diffusion signal as a sum of compartments representing varying tissue geometries (Zhang et al., 2012). However, some assumptions of the NODDI model are likely not valid in practice. The tortuosity assumption that links the extra-cellular perpendicular diffusivity to the axon volume fraction does not hold for tight axon packings (Jelescu et al., 2016). The NODDI model also sets the intra-axonal parallel diffusivity equal to the extra-axonal parallel diffusivity. However, the intra-axonal parallel diffusivity has been shown to be much higher than the extra-axonal parallel diffusivity (Jelescu et al., 2016; Novikov et al., 2018). While we altered the fixed NODDI diffusivity values to reflect what has been found in gray matter, the modelling degeneracies mentioned here could influence the NODDI metrics, such that they may not correspond to the biophysical reality of the tissue. To this end, there has been recent work which has sought to re-structure the basic microstructural building blocks used for white matter modelling to improve characterization of the gray matter. Two primary examples of this include a model which accounts specifically for the presence of the soma (Palombo et al., 2020) and a model which considers the water exchange across the cell membranes (i.e., from neurites to extra-cellular space) (Jelescu et al., 2022). Future work should look to apply gray matter specific models in the hippocampus.

Finally, the T1w/T2w ratio as a proxy for myelin content was used in the current study based on previous work that demonstrated good correspondence with cortical patterns of myelin distribution (Glasser & Van Essen 2011). However, recently it has been suggested that the T1w/T2w ratio is a suboptimal proxy of myelin in the subcortex (Arshad et al., 2017). At present, it is unclear whether the T1w/T2w ratio is also a suboptimal marker of myelin for archicortical areas such as the hippocampus. Future work should look to investigate the use of other techniques derived from quantitative MRI including magnetization transfer and myelin water imaging to get an improved measure of myelin content within the hippocampus (Tardif et al., 2016).

## 5. Conclusion

In the current study we show distinct in vivo microstructural distributions and orientations within and across the hippocampal subfields, something that has not been investigated with comparable granularity up to this point. Furthermore, we provide context for the use of surface-based approaches to investigate hippocampal microstructure.

Our findings have several important implications for future work. The hippocampus is particularly vulnerable to certain neurological diseases such as Alzheimer ’s disease and epilepsy, in which it is often one of the earliest aberrant structures (Dhikav et al., 2012). Examining the microstructure of the hippocampus at fine spatial resolutions in the simplified unfolded space, as done in this study, may provide potentially useful markers of hippocampal integrity. Furthermore, we noticed relatively large radial and tangential components of diffusion mainly in CA1 and the subiculum. Future work could attempt to tease apart these two orientationally distinct populations, providing estimates which may be useful to examine microstructurally specific deterioration. Furthermore, using the same orientation methods in this study, future work should focus on capturing multiple microstructure orientations as the hippocampus contains a complex configuration of fiber orientations. Future work could also relate all the identified OPNNMF components to demographic and cognitive variables to identify if there is a relationship between variability in cognitive performance and variability in the metrics used in this study. Finally, the microstructural metrics observed in this study appear to show good separability across hippocampal subfields, suggesting they may be sensitive to the underlying cyto- and myeloarchitectonic differences.

## Acknowledgements

This work was supported in part by funding provided by Brain Canada, in partnership with Health Canada, for the Canadian Open Neuroscience Platform initiative. BK is supported by a post-graduate scholarship from the Natural Sciences and Engineering Research Council of Canada (NSERC). JD is supported by a postdoctoral fellowship NSERC grant. ARK was supported by the Canada Research Chairs program (#950-231964), NSERC Discovery Grant (#6639), and Canada Foundation for Innovation (CFI) John R. Evans Leaders Fund project (#37427), the Canada First Research Excellence Fund, and Brain Canada. ARK and SK were supported by a Canadian Institute for Health Research grant (CIHR Project grant #366062). SK is supported by a NSERC discovery grant (#05770). This research was enabled in part by the support provided by the Digital Research Alliance of Canada. Data were provided [in part] by the Human Connectome Project, WU-Minn Consortium (Principal Investigators: David Van Essen and Kamil Ugurbil; 1U54MH091657) funded by the 16 NIH Institutes and Centers that support the NIH Blueprint for Neuroscience Research; and by the McDonnell Center for Systems Neuroscience at Washington University.

## Data and code availability

The *HippUnfold* software is openly available and can be found at https://github.com/khanlab/hippunfold. Code used for the study can be found at https://github.com/Bradley-Karat/Hippo_Macro_Micro.

Data were provided [in part] by the Human Connectome Project, WU-Minn Consortium (Principal Investigators: David Van Essen and Kamil Ugurbil; 1U54MH091657) funded by the 16 NIH Institutes and Centers that support the NIH Blueprint for Neuroscience Research; and by the McDonnell Center for Systems Neuroscience at Washington University; https://www.humanconnectome.org/study/hcp-young-adult/document/1200-subjec ts-data-release

