## Supplementary figures for "Mapping the Macrostructure and Microstructure of the in vivo Human Hippocampus using Diffusion MRI"

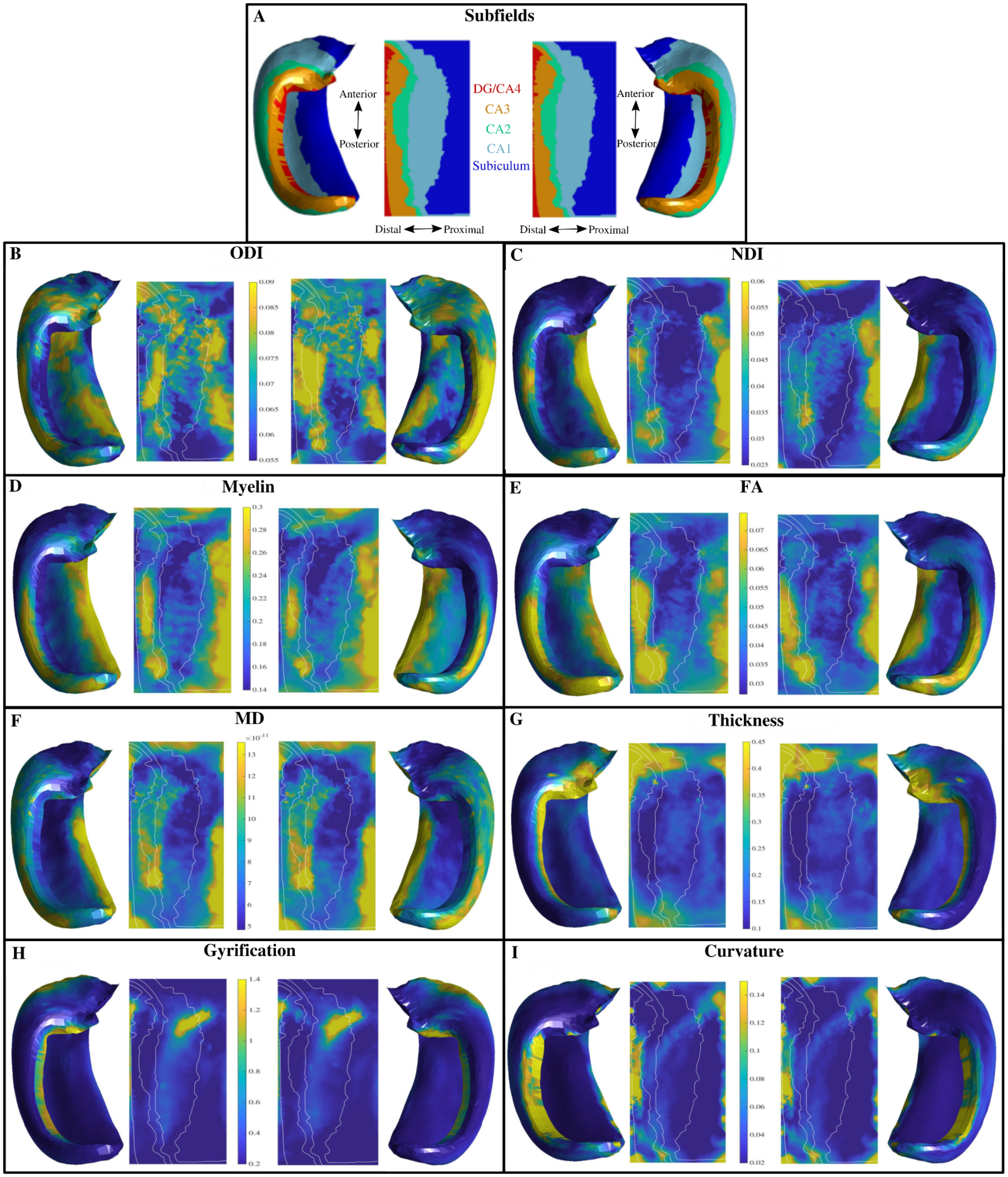


**Supplementary Figure 1.** Plots of the standard deviations for macro- and microstructure metrics on averaged hippocampal midthickness surfaces in folded and unfolded space for left and right hemispheres. (A) Left and right hippocampal subfields from a manual segmentation of a histological reference (Ammunts et al., 2013; DeKraker et al., 2020). Unfolded space is shown in the same orientation for left and right hemispheres. DG - Dentate Gyrus, CA - Cornu Ammonis.

(B,C) Orientation Dispersion Index (ODI) and Neurite Density Index (NDI) from NODDI. White lines represent subfield borders shown in (A). (D) Myelin content. (E,F) Diffusion Tensor Imaging metrics of Fractional Anisotropy (FA) and Mean Diffusivity (MD). (G-I) Macrostructure measures of thickness, gyrification, and curvature.

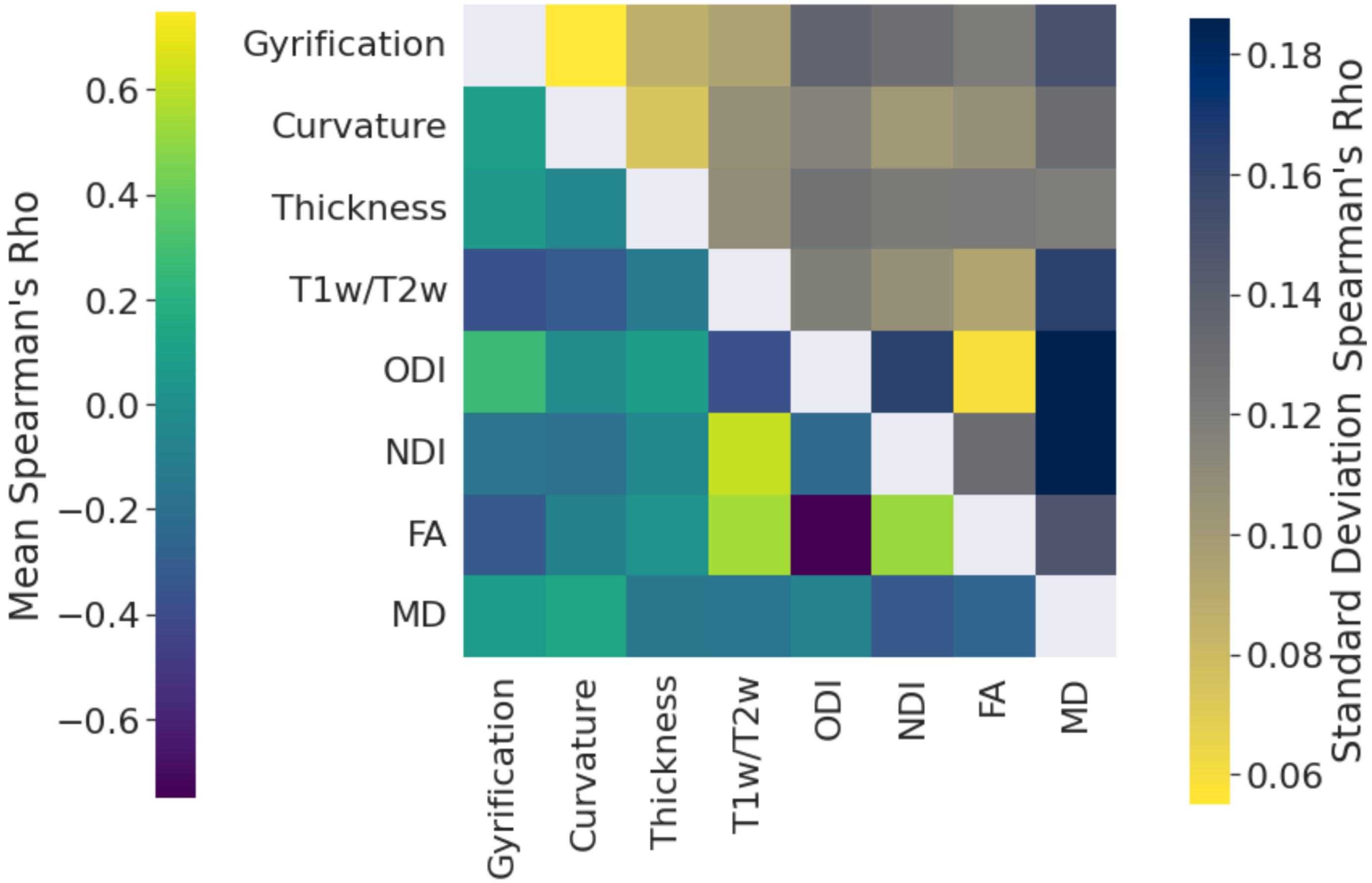


**Supplementary figure 2.** Mean and standard deviation of Spearman's rho correlations between all maps at the subject level. 200 Spearman's rho correlation values were obtained between any two maps (left and right hemisphere combined).

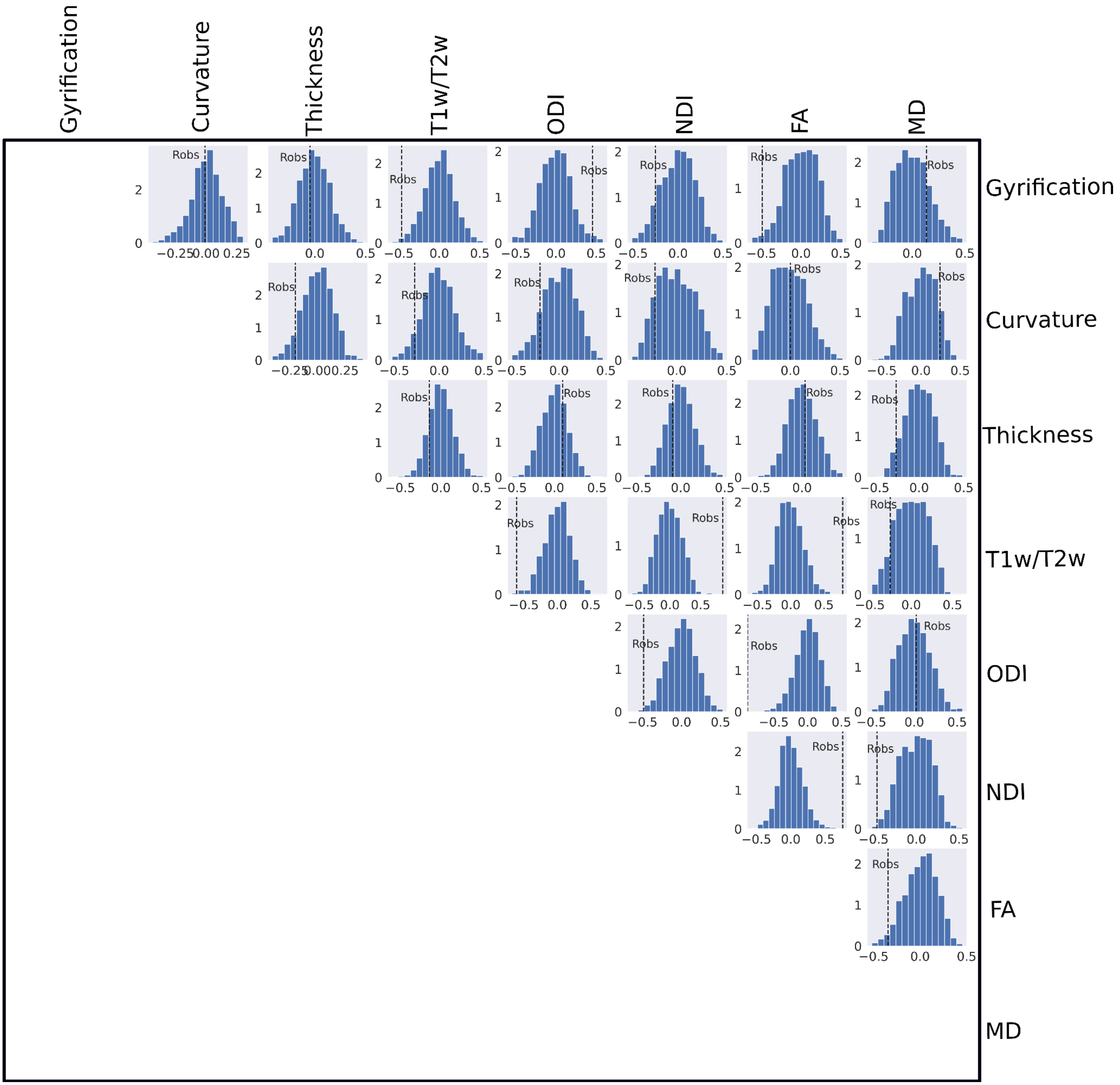


**Supplementary figure 3.** Null distributions calculated between averaged metric maps using the developed hippocampus spin test. The x-axis of each figure is the Spearman's rho value, and the y-axis is the density. The black dotted lines represent the observed association between the two maps.

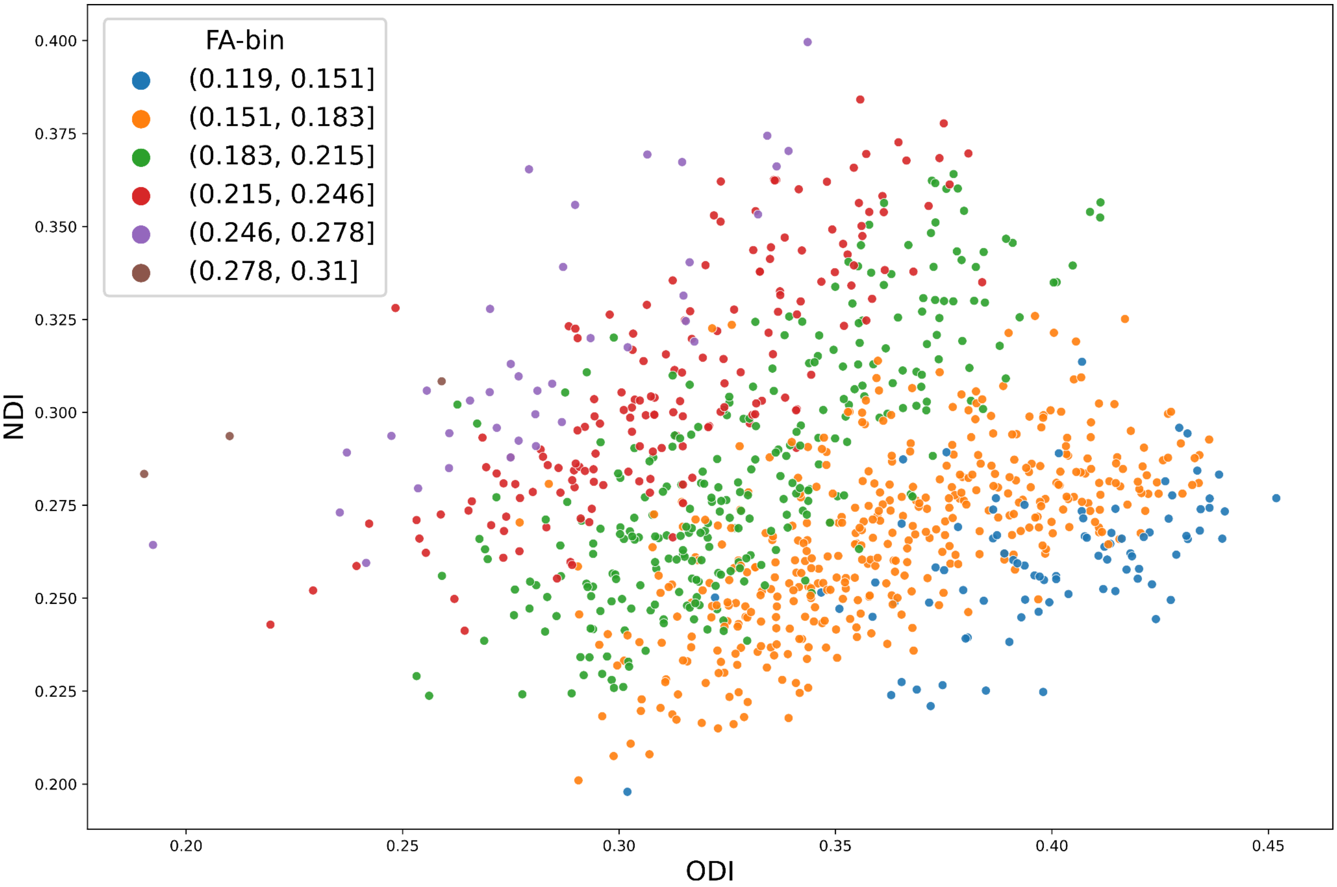


**Supplementary Figure 4.** Correlation between ODI and NDI grouped by ranges of FA values.

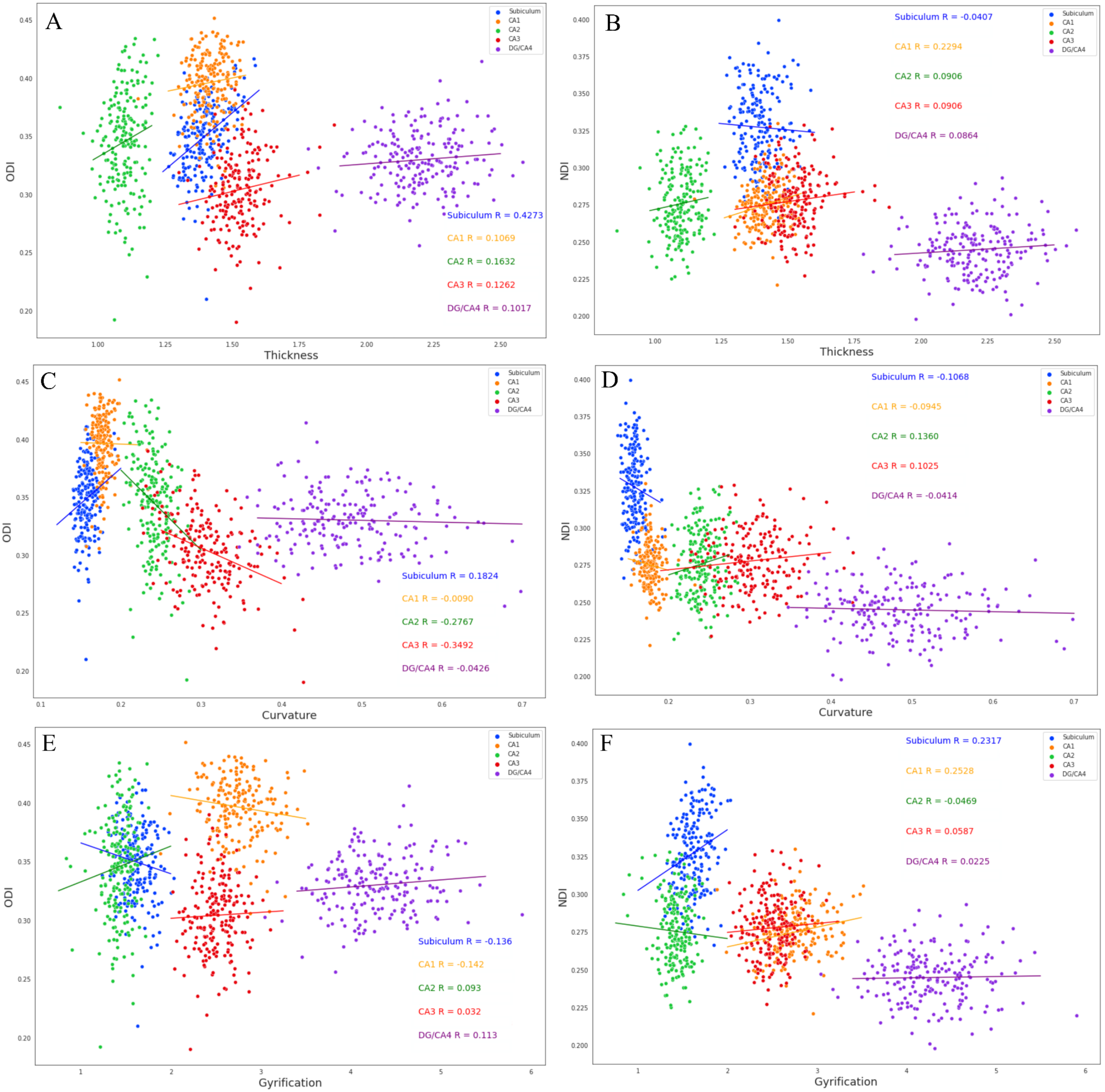


**Supplementary Figure 5.** Subfield-averaged scatter plots between macro- and microstructural measures. Linear regression was performed within each subfield to draw lines of best fit, and the correlation between macro- and microstructure was performed within each subfield.

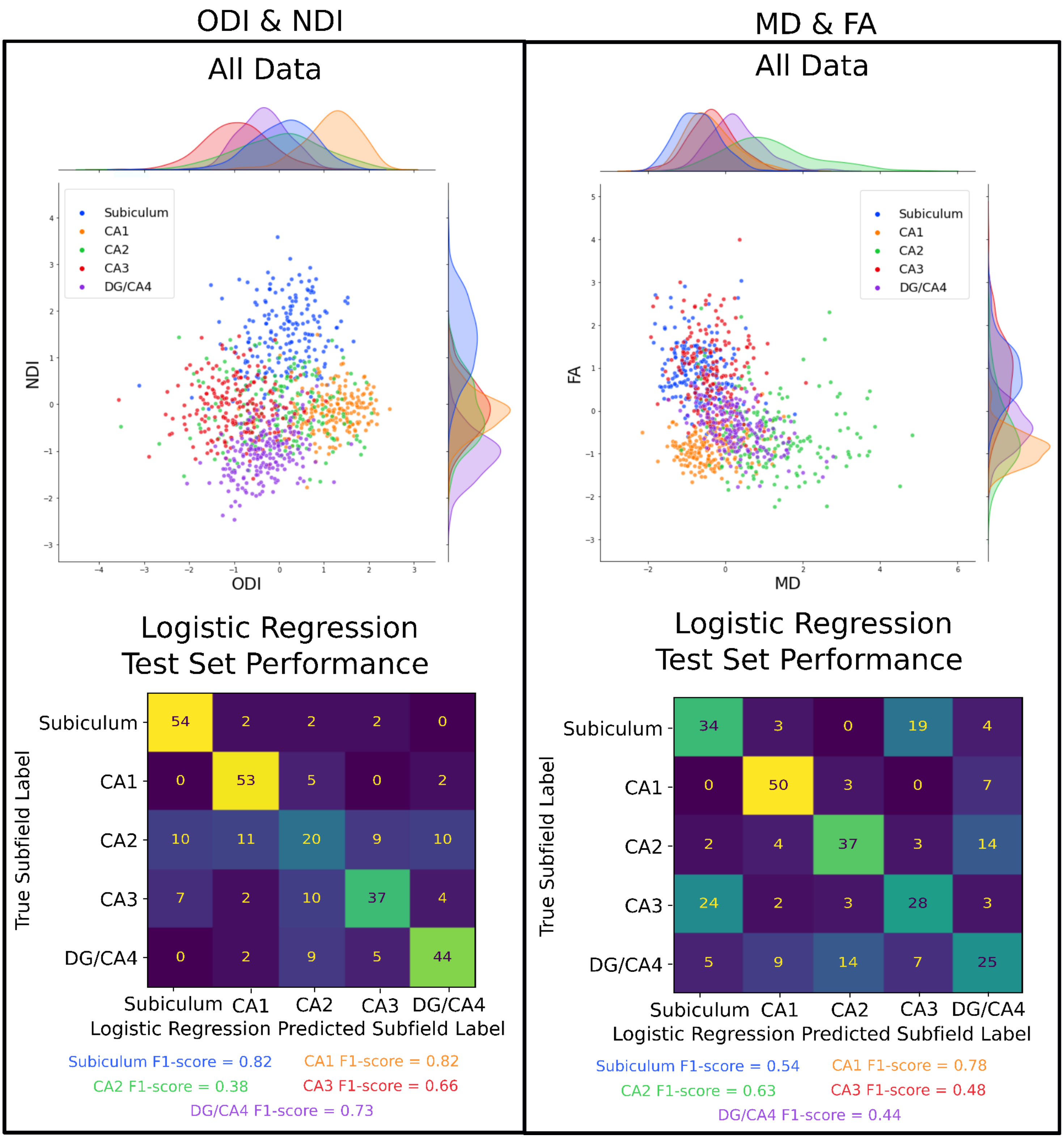


**Supplementary figure 6.** Plotting the z-scored subfield averaged measures of ODI and NDI (left) and MD and FA (right), along with the logistic regression performance on the test set for each subfield.

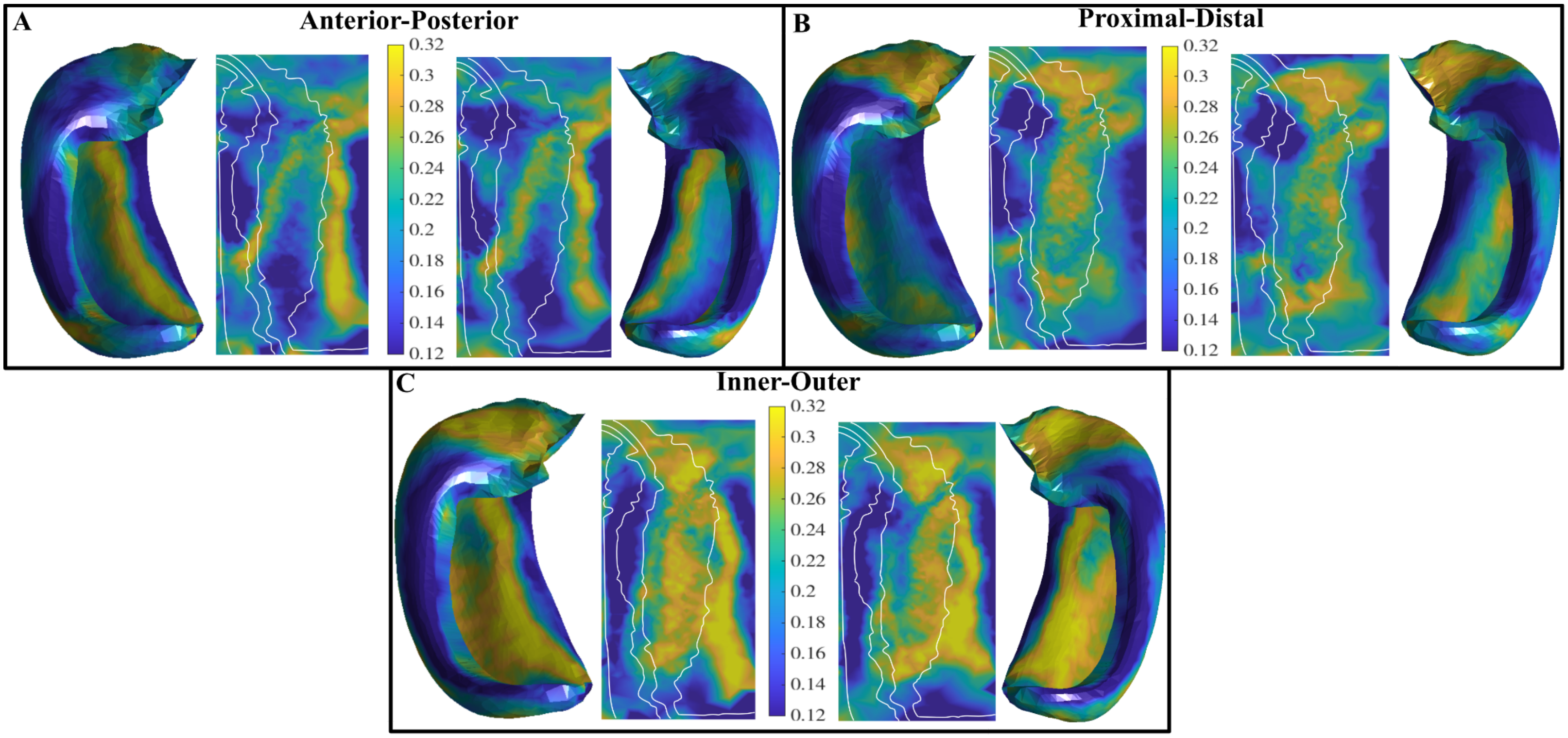


**Supplementary Figure 7.** Standard deviation of the cosine similarities between hippocampal axis vectors and NODDI vectors. Cosine similarities were sampled across the midthickness surface and are plotted on averaged surfaces.

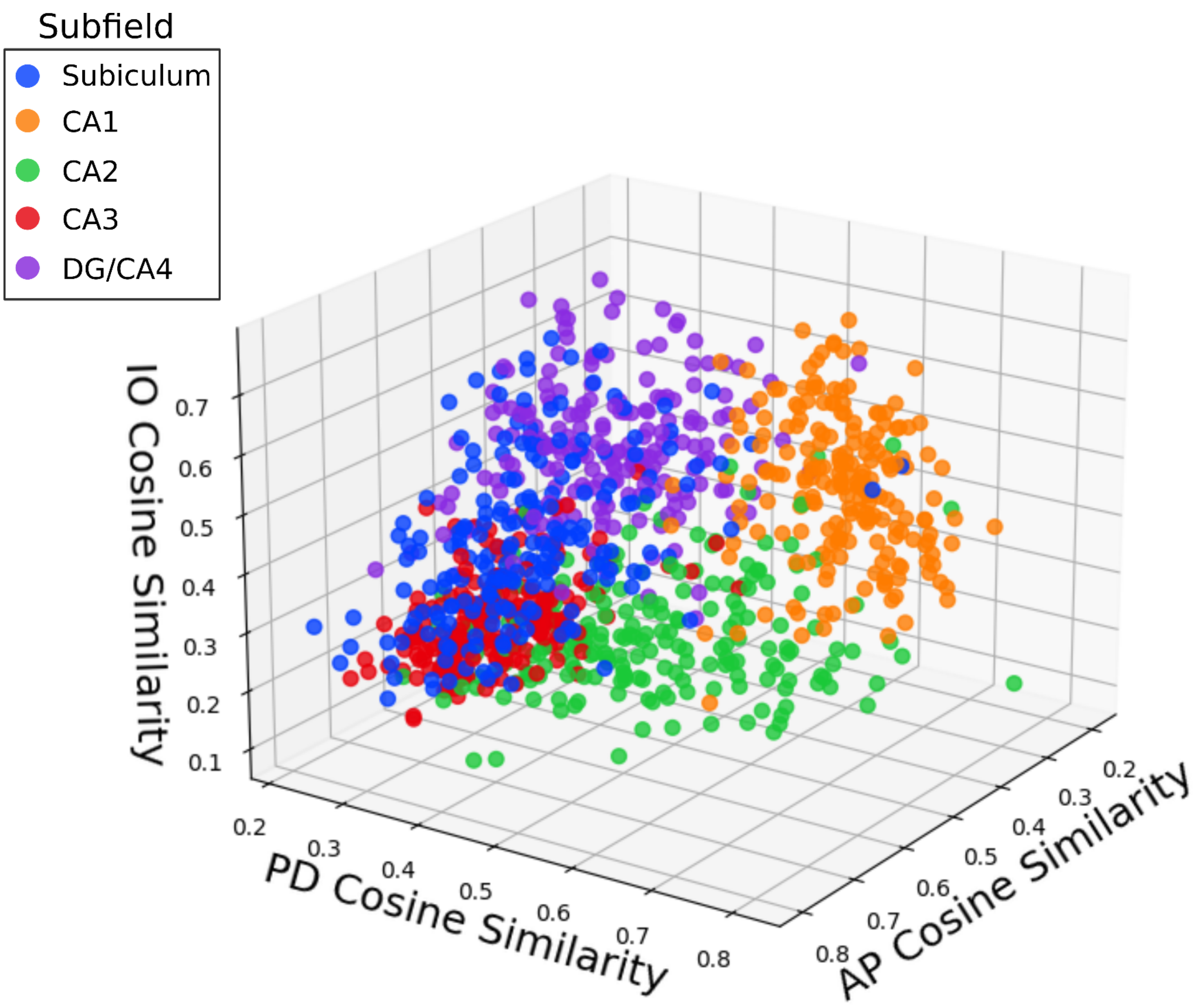


**Supplementary Figure 8.** Plotting the subfield averaged cosine similarities colour coded by subfield class.

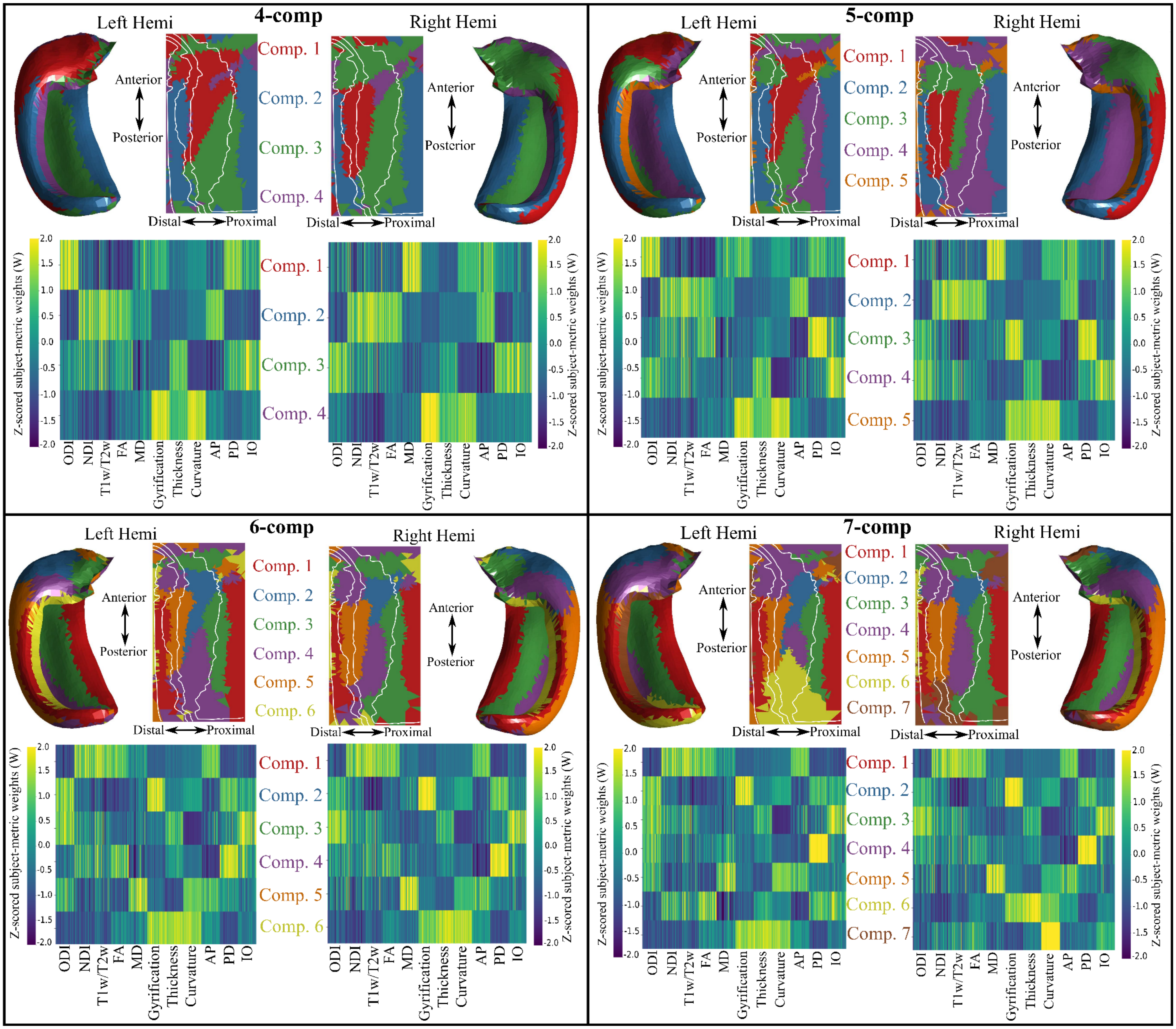


**Supplementary figure 9.** Varying the component value using all metrics for the NMF solution on the midthickness surface for left and right hippocampi. Winner-take all output at each vertex shown in folded and unfolded space at the top row of each box. White lines denote subfield borders. Bottom row in each box denotes the z-scored contribution of each metric for each component. (A) 4-component solution. (B) 5-component solution. (C) 6-component solution. (D) 7-component solution.

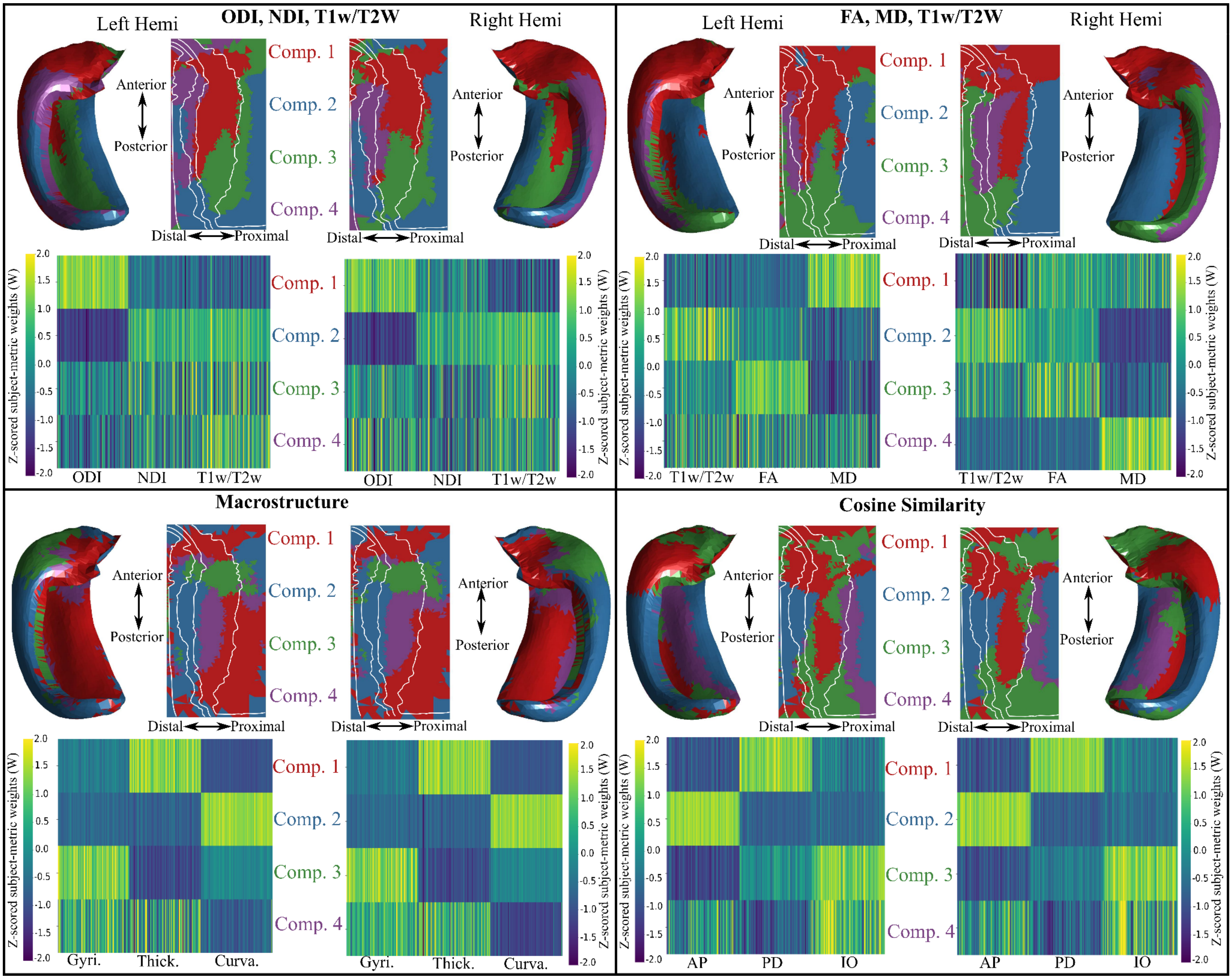


**Supplementary figure 10.** Varying the input metrics for the NMF solution on the midthickness surface for left and right hippocampi using a 4-component solution. Winner-take all output at each vertex is shown in folded and unfolded space at the top row of each box. White lines denote subfield borders. Bottom row in each box denotes the z-scored contribution of each metric for each component. (A) ODI, NDI, myelin input matrix. (B) FA, MD, myelin input matrix. (C) Macrostructure (gyrification, thickness, and curvature) input matrix. (D) Cosine similarity (AP, PD, and IO) input matrix.

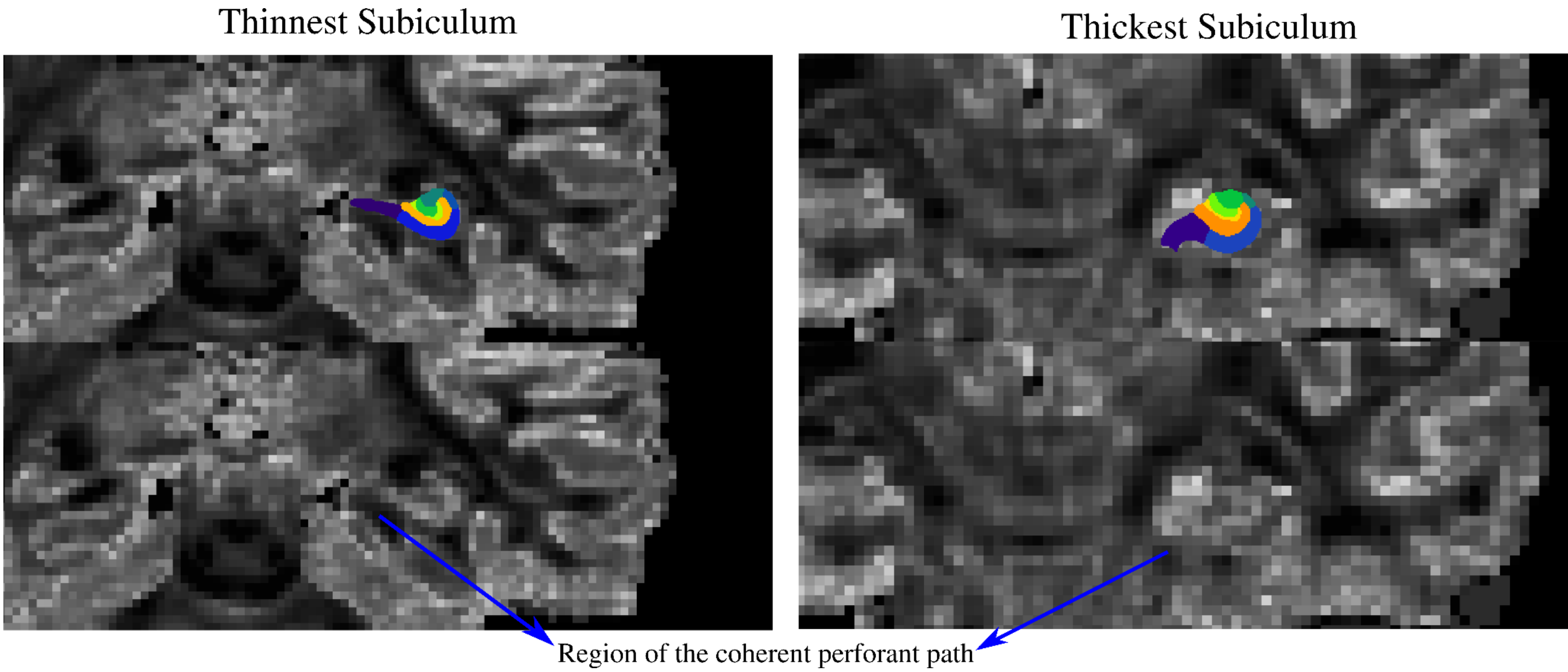


**Supplementary Figure 11.** Depicting the thinnest and thickest subiculum (purple subfield label) out of all 100 subjects plotted on top of the native space ODI image. Blue arrows point to regions of low dispersion which correspond to the highly coherent perforant path/angular bundle. Partial voluming can be seen with the thinnest subiculum, as lower ODI values from the perforant path/angular bundle are present in the gray matter. The thickest subiculum shows less partial voluming.
