## Supplementary document 1 for "Mapping the Macrostructure and Microstructure of the in vivo Human Hippocampus using Diffusion MRI"

[1]Robarts Research Institute, Schulich School of Medicine and Dentistry, University of Western Ontario, Canada, [2]Neuroscience Graduate Program, University of Western Ontario, Canada, [3]Montreal Neurological Institute, McGill University, Montreal, Quebec, Canada, [4]University Health Network, Toronto, Ontario, Canada, [5]Department of Psychology, University of Western Ontario, Canada, [6]Western Institute for Neuroscience, University of Western Ontario, Canada,

*Page 2 contains a more theoretical background for the spin test as seen in Alexander-Bloch et al., 2019. Page 3 onwards contains a more pragmatic description with additional analyses.

For our defined "plane test" we adopt the general nomenclature from the developed spin test from Alexander-Bloch et al., (2019).

### 1 Describing permutation testing in unfolded space

Let $\mathbb{P}^2$ denote a plane embedded in $\mathbb{R}^3$. Let A(x,y), B(u,v) $\in \mathcal{F}(\mathbb{P}^2)$ be two continous functions on the plane. The values of the function depend on the in-plane coordinates (x,y and u,v). These two planes represent two spatial maps of the unfolded hippocampus (see DeKraker et al., 2022). As in Alexander-Bloch et al., 2019, we assume that the anatomical alignment of these two maps is unknown. Thus there exists some set of translations and rotations T $\in$ SE(2) such that B(u,v) * T = B(x,y), that is, the transformation anatomically aligns the two planes. SE(2) is the set of all rigid translations and rotations in 2D euclidean space. There also exists some measure of association ($\psi$), between maps A and B that maps to the space of real numbers. The spatial permutation test then looks to statistically examine whether two surfaces are correlated. As in Alexander-Bloch et al., (2019), the null hypothesis states that the values on the surface maps does not provide information about their alignment:

$$H_0 : T|A(x,y), B(u,v)\ UniformSE(2) \tag{1}$$

That is, given two maps A and B, any transformation from the set SE(2) leads to approximately the same result (no gain in confidence of any particular alignment of the maps given by T). Let $R_{obs}$ denote the association given by ($\psi$) in their observed alignment and let $P_0$ denote the uniform distribution on SE(2). The p-value is calculated as:

$$p = \int_{SE(2)} I(\psi(T) \geq \psi(R_{obs}))dP_0(T) \tag{2}$$

and the null hypothesis is rejected if p < $\alpha$ for some predetermined threshold. See theorem 1.1 in Alexander-Bloch et al., (2019) for justification on the following: p = P($\psi$(R) $\geq \psi(R_{obs})$), that is, the p-value is the number of permuted association values that are greater then or equal to the observed association value, divided by the total number of permuted values. Given the set of all T applied to a plane, we generate a spatially permuted null distribution of $\psi$ values. We then compare our observed $\psi$ when the surfaces are anatomically aligned to this null distribution. The null hypothesis is then

$H_0$: The observed association given by $\psi$ between A and B provides no information about how the surfaces are anatomically aligned

To reject the null hypothesis is to say that the extent of the association when the surfaces are aligned is improbable to have resulted from a random alignment of the surfaces.

The goal of the spin test is to compute spatial autocorrelation (SA) corrected p-values between two brain maps. Contemporary cortical spin-testing works by rotating a spherical representation of a cortical map. However, the geometry of the hippocampus is not conducive to a spherical representation. Thus, we are looking to adapt the spin test to the HippUnfold planar representation. The code for the spin testing can be found here: [https://github.com/Bradley-Karat/Hippo_Spin_Testing](https://github.com/Bradley-Karat/Hippo_Spin_Testing)

The vertex spacing of the hippocampal surfaces calculated via HippUnfold can be represented by vertex spacings of 0.5mm, 1mm, and 2mm. The first step of the spin test is to interpolate these non-uniform vertex spacings in unfolded space (see figure S1) to an evenly spaced grid (defined as unfold-iso within HippUnfold). The evenly spaced unfolded grid allows us to capitalize on previous functions made for 2D image processing (like .jpg and .png images).

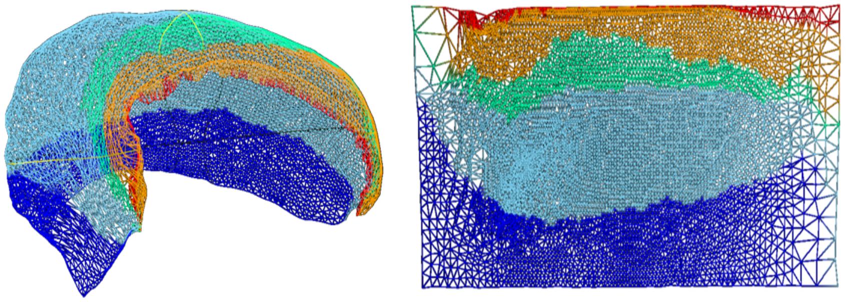

**Figure S1.** Vertex spacing of 1mm in folded and unfolded space. The vertex spacing is defined such that it is approximately uniform in folded space (left), which leads to stark non-uniformities in unfolded space (right).

To transform our plane, the spin test uses the rotate and shift functions from Scipy (https://scipy.org/) with wrapping to define our boundary conditions. An example of what these transformations look like can be seen in figure S2. This can be related to transformation on a torus geometry, where the anterior connects to the posterior and the subiculum connects to the dentate gyrus.

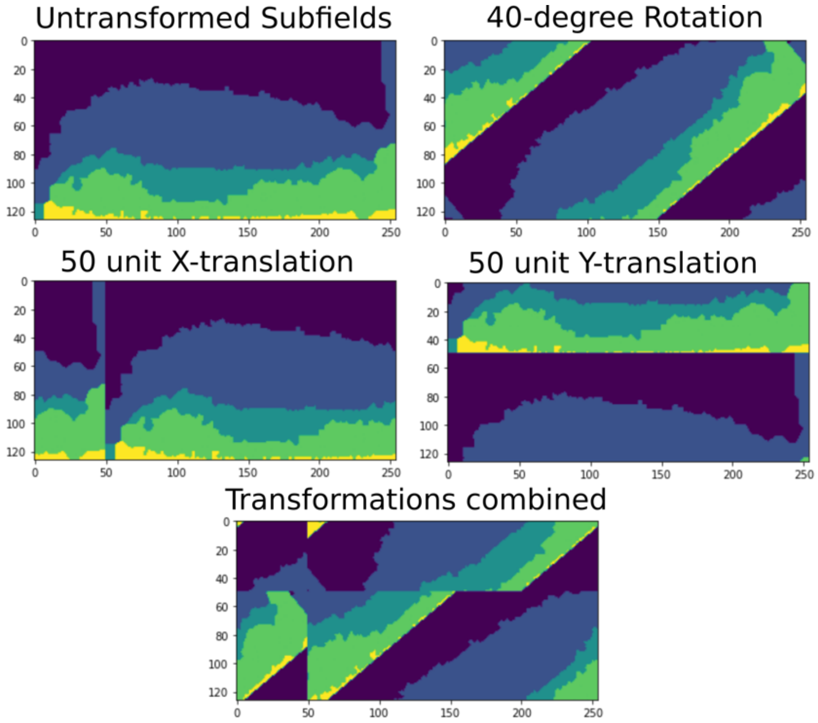


**Figure S2.** An example showing transformation behaviour on the unfolded surface representation of the subfields. The “wrap” boundary condition results in vertices that “fall” off the edge to re-appear in the same location just on the opposite end of the plane. For example, if we consider an X-translation, the Y-coordinate of all vertices remains constant even after encountering a boundary.

After applying a random rotation, X-translation, and Y-translation, we can compare the untransformed map with the transformed map with any metric of choice. This will most commonly use Pearson's R or Spearman's Rho. We can then repeat this process with a new random transformation to build up a null distribution of our metric of choice. An example of this can be seen in figure S3. We can then define the p-value as the number of permuted associations that were greater than or equal to our observed association, divide by the total number of permutations.

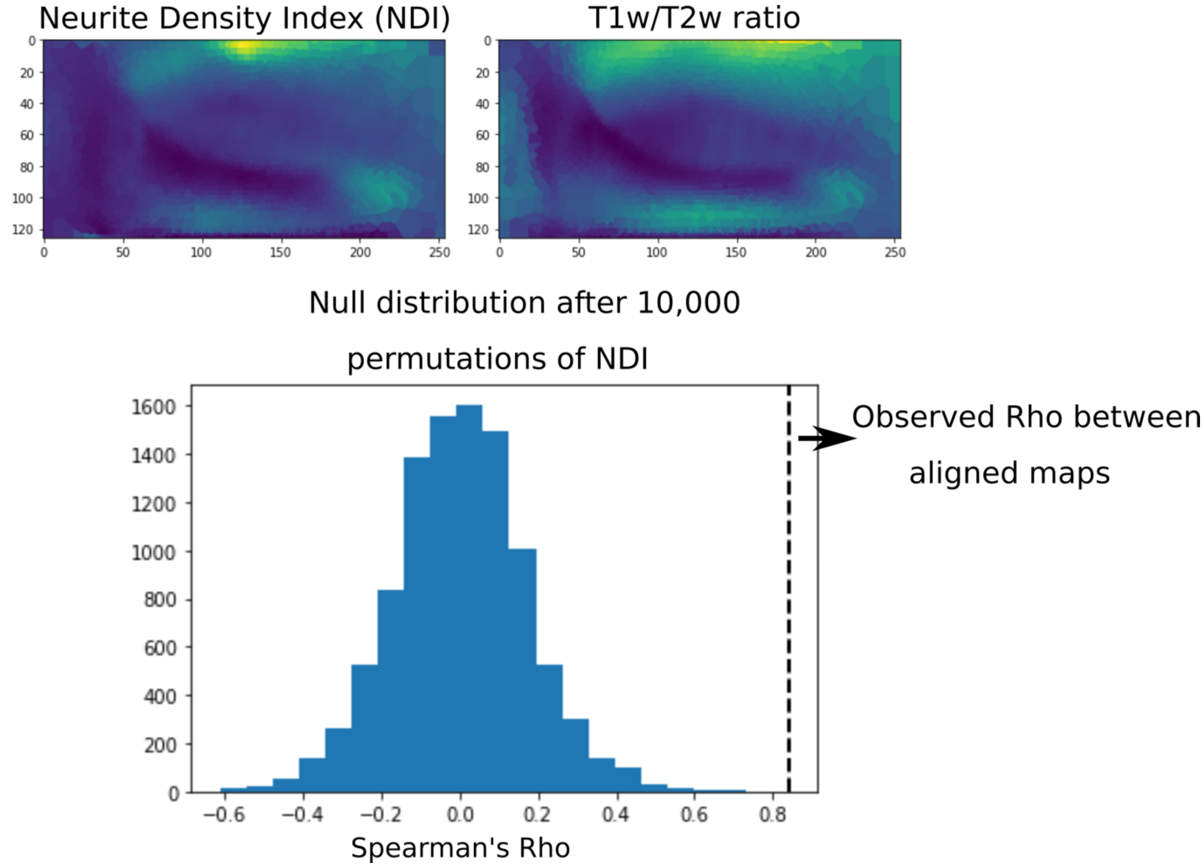


**Figure S3.** Top row shows two different continuous maps in unfolded space that we will correlate. The bottom histogram shows the null distribution obtained after applying the spin test to the NDI map for 10,000 permutations. The dotted line represents the observed correlation between the two anatomically aligned maps (top row). In this case, we would reject the null hypothesis that there is no anatomical correspondence between these maps.

### Type 1 error rate analysis

We performed tests to assess whether our implementation of the spin test controls the type 1 error rate. We used a set of continuous maps sampled on the hippocampal midthickness surface seen in figures 2 and 4 of the main text. We sought to generate maps under the null hypothesis. This means that we wanted two maps where their observed association provided no information about their anatomical alignment, or that the observed association could have plausibly resulted from a random alignment. To generate maps under this hypothesis, we took a subset of 4 maps that remained fixed, and a different subset of 5 maps which were transformed randomly. Randomly transforming the subset of 5 maps before permutation testing removed their anatomical alignment with the fixed maps, such that the observed association was a result of a random alignment. The null hypothesis in this case should then be true, since the observed association was given by a random alignment. We then performed the spin test as usual between all fixed and rotated maps to generate a null distribution of correlation values and a p-value using 2000 permutations. This process of applying a random transformation and then subsequent spin-testing was repeated 500 times for each map. That is, within each comparison of a fixed to transformed map, we generated 500 p-values. We then determined the number of p-values that were below our pre-specified thresholds of 0.05 and 0.01 for significance, which is the type 1 error rate. If the type 1 error rate was controlled, we would expect around 5% or 1%, respectively, of all our p-values to be significant. The fixed maps were FA, MD, AP cosine similarity, and IO cosine similarity. The rotated maps were ODI, NDI, T1w/T2w, gyrification, and PD cosine similarity as seen in figures 2 and 4.

At $\alpha = 0.05$ around 5% of the randomly transformed data resulted in false positives, and at $\alpha = 0.01$ around 1% of the randomly transformed data resulted in false positives, thus it appeared that the type 1 error rate was controlled.

**Table S1.** Type 1 error rate analysis of the hippocampus spin test.

| Fixed map | Mean (SD) at $\alpha = 0.05$ | Mean (SD) at $\alpha = 0.01$ |
|---|---|---|
| FA | 0.052 (0.019) | 0.008 (0.007) |
| MD | 0.052 (0.020) | 0.01 (0.006) |
| AP cosine Similarity | 0.057 (0.019) | 0.013 (0.007) |
| IO cosine Similarity | 0.064 (0.020) | 0.012 (0.007) |

### Variogram Method Comparison

We then looked to compare our developed spin test method against the variogram method (Burt et al., 2020) which works by generating surrogate maps which have SA matched to the SA of a target brain map. This method works on subcortical and volumetric data, and thus generalizes beyond the cortical spin test which requires a spherical representation. We used the brainstat (https://brainstat.readthedocs.io/en/master/index.html) implementation of the variogram method. In figure S4 we show the unfolded space variogram and our spin test null distribution for a group of fixed maps (ODI, MD, and NDI) and a group of permuted maps (gyrification, FA, and T1w/T2W). In some cases, we can see that the spin test null distribution has a lower variance than the variogram method. However, we can see that they are still approximate to each other, and the derived conclusions (i.e. the significance of the correlation) is equal in all cases for both methods. An important note is the computation time differences. It took approximately 30-40 minutes to generate 1000 surrogate maps using 12 CPUs (without subsampling) with the variogram method to then build our null distribution. In contrast, our spin test implementation took approximately 10-20 seconds to generate 1000 permutations to build our null distribution using the same computational resources.

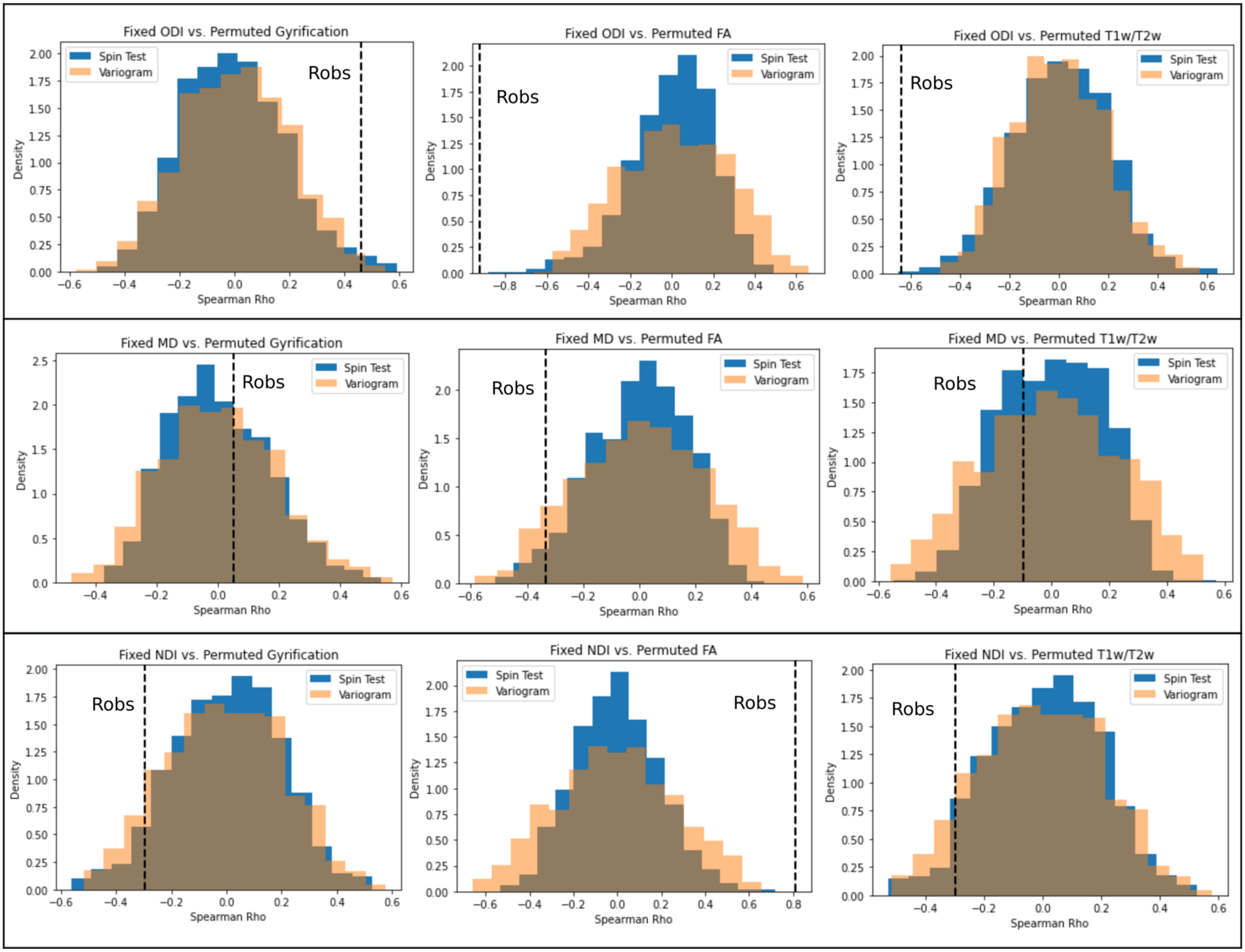


**Figure S4.** Variogram matching method vs. spin test comparison for multiple fixed and permuted maps.
